# Super-Resolution Atlas of SARS-CoV-2 Infection Reveals Protease-Dependent Organelle Maturation, dsRNA Landscapes, and Intracellular Structural Proteins

**DOI:** 10.1101/2025.08.15.670620

**Authors:** Leonid Andronov, Mengting Han, Ashwin Balaji, Yanyu Zhu, Lei S. Qi, W. E. Moerner

## Abstract

Coronaviral replication depends on double-membrane vesicles (DMVs), yet where polyprotein processing occurs and how replication organelles mature remain unresolved. We built a multi­color super-resolution atlas of SARS-CoV-2 RNA, non-structural and structural proteins in infected human cells, combining 3D single-molecule localization with radial and angular pair-correlation analysis. The atlas uncovers fundamental spatial principles. First, nsp5 (3CL^pro^) localizes within the DMV lumen near pores, with nsp7–nsp16 positioned interior to nsp4, supporting a protease-dependent maturation model in which nsp5 action permits membrane closure and subsequently completes intravesicular polyprotein processing to activate replication complexes. Second, we identify dsRNA connectors bridging DMVs and non-canonical, nsp3/4-lacking dsRNA granules decorated with replicase components, consistent with a condensate-mediated route for trafficking replication intermediates. Third, mapping structural proteins reveals M- and S-positive virion assembly intermediates along the secretory route, while premature intracellular S1 shedding captures the baseline instability of the original virus isolate prior to evolutionary adaptation. Finally, the antiviral nirmatrelvir induces multilayered bodies of uncleaved polyproteins (nsp4–5–10–16) that persist after washout and precede rapid rebound, suggesting a drug-induced reservoir state. Beyond mapping the viral architecture, this atlas resolves the spatial context of proteolysis and organelle remodeling, providing a framework for pan-coronaviral mechanisms and antiviral design.

## Introduction

Positive-strand RNA viruses replicate in membrane-bound organelles, yet where coronaviral polyproteins are processed and how replication organelles mature remain unclear. The SARS-CoV-2 viral RNA molecule codes for 29 proteins, whose expression yields polyprotein structures and modifies endoplasmic reticulum (ER) membranes in infected cells (Fig. 1A). The intracellular localization of several SARS-CoV-2 proteins has been investigated using diffraction-limited (DL) microscopy in infected cells^1,2,3^; however, due to the diffraction limit (∼200 nm), DL microscopy cannot resolve key virus-induced structures, such as double-membrane vesicles (DMVs) or virions, nor can it distinguish tiny viral RNA structures.

**Fig. 1.**
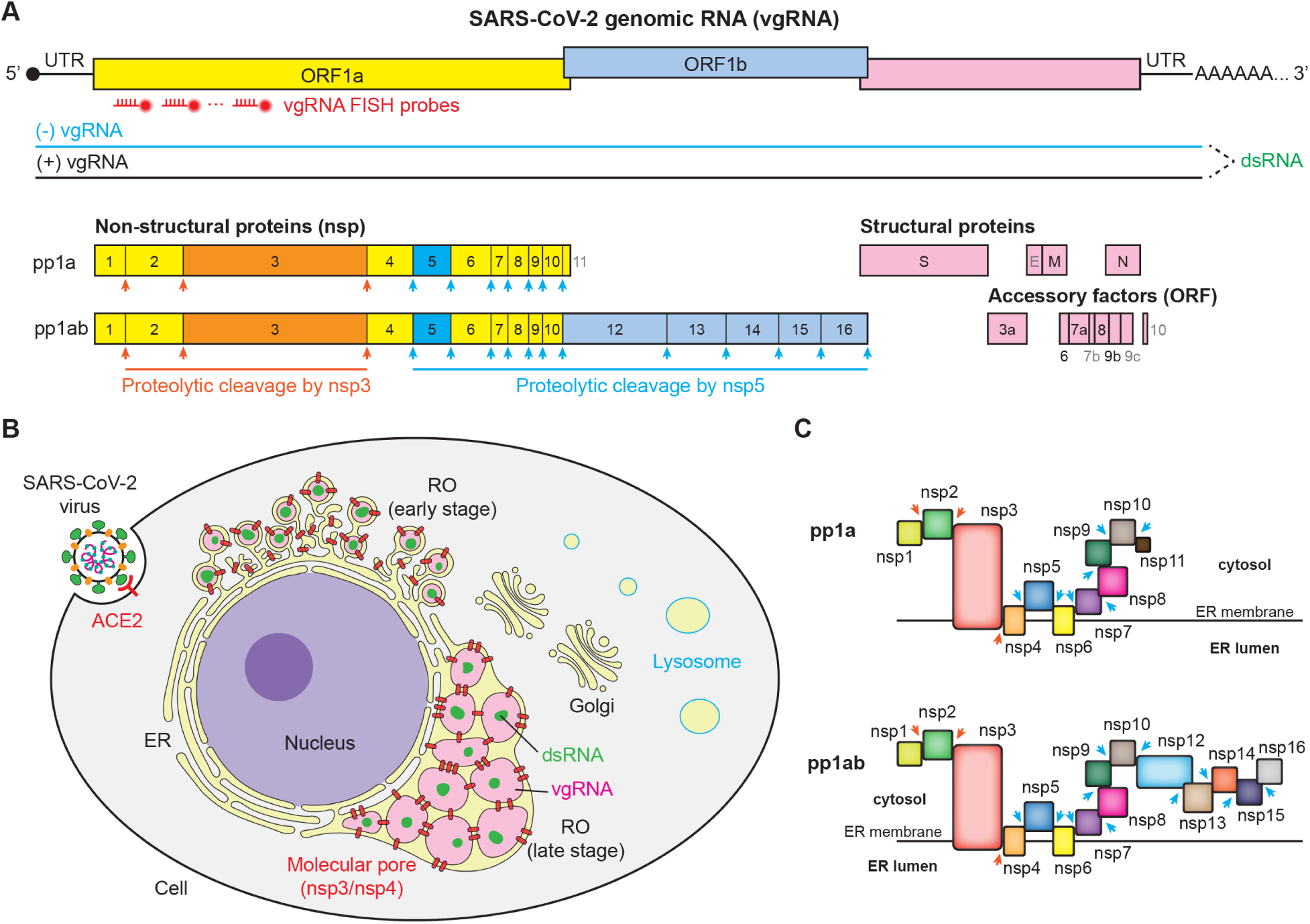
SARS-CoV-2 genome organization, replication organelles, and polyprotein topology. (A) Schematic of the SARS-CoV-2 genome. Targets explicitly labeled and investigated in this study are indicated in black text, while uninvestigated regions are shown in gray. Red markers indicate the binding sites for the vgRNA FISH probes. (B) Model of a SARS-CoV-2–infected A549-ACE2 cell highlighting the structural evolution of viral replication organelles (ROs) and their previously reported intracellular localizations. Key cellular targets investigated in this study are indicated. (C) Membrane topology of the uncleaved viral polyproteins pp1a and pp1ab. The schematic illustrates the relative positions of the individual non-structural protein (nsp) domains with respect to the endoplasmic reticulum (ER) membrane following translation. Arrows indicate known proteolytic cleavage sites targeted by nsp3 (orange) and nsp5 (blue).

Coronaviruses remodel ER membranes into DMVs with narrow pores that connect the lumen to the cytoplasm, and replication has been linked to these compartments^4^. By sequestering immunogenic intermediates (dsRNA, uncapped RNA), DMVs likely serve as the site of SARS-CoV-2 genome replication, consistent with accumulating evidence^5,6,7^. While foundational immune-EM studies have long placed replication-transcription complexes (RTCs) within the DMV lumen^8^, the mechanism by which these large proteins enter the sealed DMV has remained highly elusive^9^. It has been recently hypothesized that RTCs might be incorporated during DMV formation^9,10,11^, yet studies still conflict on the precise architecture, debating whether specific components reside inside the lumen^5,10^ or outside on the cytoplasmic face^12^.

Super-resolution (SR) microscopy localizes molecules with a resolution of tens of nanometers using blinking single fluorophores, collectively known as Single-Molecule Localization Microscopy (SMLM)^13^, including Stochastic Optical Reconstruction Microscopy (STORM)^14^. One related study combined focused ion beam–scanning electron microscopy (FIB-SEM) with Stimulated Emission Depletion (STED) to examine infection-induced intermediate-filament remodeling^15^. While electron microscopy (EM) visualizes membranes well, proteins and nucleic acids are less readily identified. Previously, we applied SR with immunofluorescence (IF) and RNA fluorescence in situ hybridization (FISH) to SARS-CoV-2–infected cells, mapping vgRNA, dsRNA, RdRp, and nsp3 within replication organelles (ROs)^5^. Those data established perinuclear, ER-delimited ROs containing nsp3/nsp4 pores and central dsRNA/vgRNA (Fig. 1B). However, the protease-associated maturation of replication organelles, a process central to antiviral discovery, remained unresolved.

Translation of the vgRNA produces polyproteins containing nsp1–nsp16^16^, several of which are membrane-bound (Fig. 1C). What remains unsolved is how polyprotein processing positions the constituent NSPs in infected cells, particularly which proteins reside inside versus outside DMVs and how this spatial arrangement relates to replication.

To address this, we labeled and imaged nearly all SARS-CoV-2 non-structural and structural proteins (excluding only nsp11 and E due to antibody limitations; gray in Fig. 1A) along with dsRNA and vgRNA in A549 human lung epithelial cells using 2D and 3D single-molecule SR microscopy. Where possible, imaging was three-color, yielding a quantitative spatial atlas that provides direct mechanistic insights into viral replication and assembly. Beyond prior diffraction-limited localizations, we identify (a) novel dsRNA connectors linking ROs; (b) intravesicular organization of the DMV-RO defined by the positioning of nsp5, the polyprotein protease, relative to the nsp3/nsp4 pore-forming complex; and (c) diverse morphologies of the (M) protein across cellular compartments in infected cells. Additionally, treatment with an nsp5 protease inhibitor, nirmatrelvir, induces multilayer, shell-like membrane bodies enriched for multiple NSPs. Together, our comprehensive super-resolution measurements offer deeper insight into the structures and mechanisms of viral infection, coupling a spatial atlas to protease-associated RO maturation and providing targets and hypotheses for antiviral intervention.

## Results

We used A549-ACE2 human lung carcinoma epithelial cells that stably express the human ACE2 receptor (angiotensin-converting enzyme 2), which provide a more relevant respiratory model than Vero E6 or HEK293T-ACE2^17^. Infected cells were imaged by super-resolution (d)STORM fluorescence microscopy^18^, which, unlike DL light microscopy or EM, offers both nanometer-scale resolution and molecular specificity^19^. For three-color SR microscopy, we simultaneously imaged two spectrally close far-red fluorophores, Alexa Fluor 647 (AF647) and CF680, separated by ratiometric demixing^20^, followed by acquisition of emission from a third, spectrally distinct fluorophore CF583R^21^. This leverages low cellular autofluorescence and high-performance dyes in the red and far-red spectral regions and reduces the number of different fluorophores imaged per frame.

Viral and host proteins (Fig. 1) were labeled by IF using a virus-inactivation-compatible protocol that preserves DMV ultrastructure^5^. We validated antibodies for 15 NSPs (nsp1–10, nsp12–16), 3 structural proteins (S/M/N), and 5 accessory proteins (ORF3a/6/7a/8/9b). Several targets (nsp1/3/4/5/7/12, N, ORF9b) had antibodies from two species (e.g., rabbit, mouse, and sheep) that allow for multiplexing using corresponding secondary antibodies labeled with spectrally distinct dyes (Supplementary Table 1). SARS-CoV-2 dsRNA was detected with the J2 monoclonal antibody^22^, as this provides high sensitivity and specificity in our experimental conditions^5,23^, and vgRNA was labeled by RNA FISH^5,24^. We also fluorescently labeled the human small ribosomal subunit 40S using RNA FISH against 18S ribosomal RNA (rRNA), which provides context for nsp1 and virus-induced DMVs. We adjusted the labeling protocol to combine RNA FISH and IF labeling in the same sample^5^ (Methods).

Non-structural proteins (NSPs, 1–16) derive from SARS-CoV-2 polyproteins pp1a/pp1ab, translated from ORF1a/ORF1ab (Fig. 1A). The pp1a/pp1ab polyproteins are cleaved by nsp3 (papain-like protease, PL^pro^) at specific sites between nsp1–nsp4, and by nsp5 (3C-like protease, 3CL^pro^, known as the main protease, M^pro^) at the remaining sites between nsp4–nsp16^25^. Unlike structural virion proteins, NSPs regulate the intracellular life cycle of the coronavirus, remodeling membranes, forming ROs/DMVs, and executing replication and transcription of the viral genome. For this reason, most NSPs are cytoplasmic and associated with ROs. While nsp3 and nsp4 localization is defined by the DMV pore structure^4^, the positioning of many other NSPs remains unclear. Here, we use nsp3/nsp4 as DMV-membrane references and dsRNA/vgRNA as DMV-interior markers^5,6,7,10^ to map all NSPs at the nanometer scale in infected cells.

Although all samples were fixed at 24 hours post-infection (hpi), the viral life cycle is asynchronous. To distinguish functional phases of replication, we classified cells into an ‘early morphological stage’ (characterized by isolated DMVs, diameter ≲ 200 nm) and a ‘late morphological stage’ (characterized by merged ROs and vesicle packets) (Supplementary Fig. 1), following the morphological transitions we previously characterized^5^ (classification criteria detailed in Methods).

### Ribosome-bound nsp1 is excluded from DMVs, and nsp2 shifts from DMV/CMs to fragmented Golgi during infection

Nsp1 inhibits host protein translation by engaging 40S ribosomes and the mRNA export receptor NXF1^26,27^. While DL studies have reported diffuse cytoplasmic nsp1 in SARS-CoV-2–infected cells^3,28,29^, its nanoscale localization was unknown. Using three-color SMLM, we co-imaged nsp1 with dsRNA (labeling the DMV interior)^5^ and nsp4 (labeling DMV pores)^4^. DMVs appeared as incomplete ring-like structures of nsp4 surrounding dsRNA clusters in the cytoplasm (Fig. 2A). Nsp1, however, showed no DMV-associated patterns and was depleted from DMV interiors, appearing diffusely in the cytoplasm outside DMVs (Fig. 2A) as isolated puncta. Because nsp1 blocks the mRNA entry channel on the 40S ribosomal subunit^26^, we labeled 18S rRNA to mark 40S subunits via FISH staining (Fig. 2B). 18S rRNA displayed a very sparse signal in the nucleus, somewhat denser punctate signal in the nucleolus and, as expected, very dense, widely scattered punctate signal in the cytoplasm, corresponding to ribosomes, polysomes, and individual 40S subunits^30^ (Fig. 2B). Nsp1 localized in the same cytoplasmic regions as 18S rRNA. To quantify 18S rRNA and nsp1 colocalization, we computed the bivariate pair-correlation function *g_12_(r)*, which measures the distribution of distances between localizations of two species; a peak at distance r_0_ indicates that the two species co-occur at that separation more frequently than expected under complete spatial randomness ^5^. For 18S rRNA and nsp1, *g_12_(r)* peaked at r = 0 nm (Fig. 2B inset), meaning that these two species tend to have a minimal distance between them, demonstrating that nsp1 indeed associates with 40S ribosomal subunits in the cytoplasm of infected cells.

**Fig. 2:**
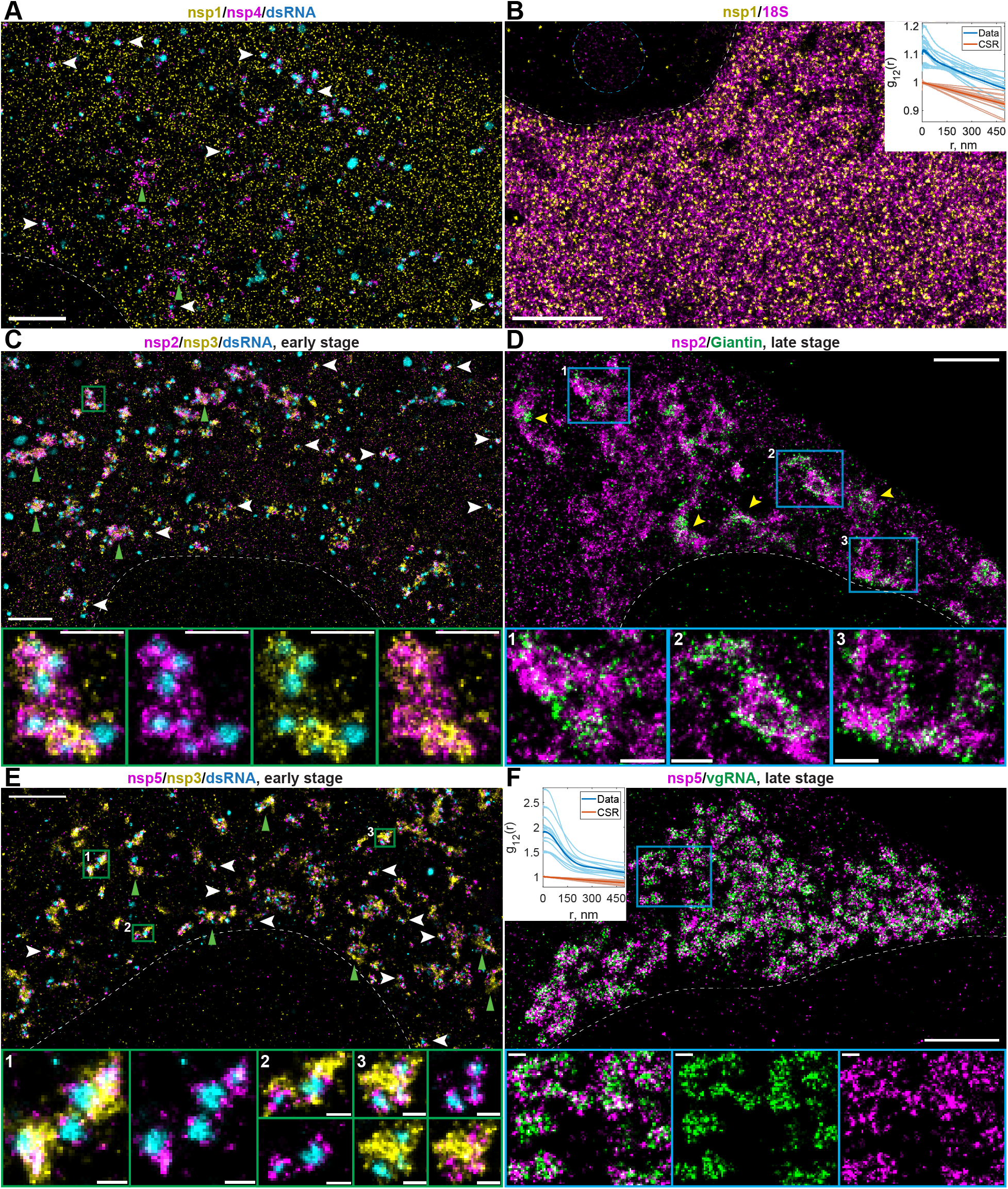
Nanoscale cellular localization of nsp1, nsp2, nsp3, nsp4, and nsp5. (A) Representative three-color SR image of a SARS-CoV-2–infected A549-ACE2 cell at an early infection stage, labeled for nsp1 (yellow), nsp4 (magenta) and dsRNA (cyan). Nsp1 localizes diffusely throughout the cytoplasm, while nsp4 encapsulates round dsRNA clusters (double-membrane vesicles, DMVs). (B) Representative two-color SR image of nsp1 (yellow) and 18S ribosomal RNA (rRNA) (magenta) indicates diffused localization of both targets within the same regions of the cytoplasm. Inset: pair-correlation functions between nsp1 and 18S localizations peak at *r* = 0 nm, suggesting an association between nsp1 and 18S rRNA. The blue dashed line approximates the position of a nucleolus, characterized by an increased density of 18S rRNA. (C) Representative SR image of an infected cell at an early infection stage, labeled for nsp2 (magenta), nsp3 (yellow) and dsRNA (cyan). Nsp2 and nsp3 co-encapsulate dsRNA clusters (DMVs) and localize between accumulations of DMVs (putative convoluted membranes, CMs). (D) In late infection, a significant portion of cellular nsp2 (magenta) localizes to fragmented and dispersed Golgi bodies (labeled with an anti-Giantin antibody, green). Bottom panels (1–3) display magnified views of selected Golgi bodies; yellow arrowheads indicate additional examples of Golgi bodies within the large image. (E) SR image of a cell in early infection suggests localization of nsp5 (magenta) within DMVs, as indicated by nsp3 (yellow, DMV membrane) and dsRNA (cyan, DMV core). (F) SR image of a cell at a late infection stage indicates that nsp5 (magenta) localizes to vgRNA-labeled (green) DMVs. Inset: pair-correlation functions between nsp5 and vgRNA localizations in late infection peak at *r* = 0 nm, indicating a strong correlation between these two targets. In the pair-correlation function plots in (B) and (F), faint blue lines represent individual cells, faint red lines represent g_12_(r) calculated assuming the complete spatial randomness (CSR), and bold lines represent the mean values across all analyzed cells. Color-framed boxes indicate magnified regions shown in the corresponding panels below. White dashed curves denote the approximate nuclear edge. White arrowheads in (A), (C), (E) indicate selected individual DMVs, whereas green arrowheads point to examples of DMV aggregates associated with nsp3-labeled CMs. Scale bars: 2 µm (large images), 500 nm (magnified images in (C), (D)), and 200 nm (magnified images in (E), (F)).

Nsp2 enhances viral infection via interaction with host proteins^31,32,33^, yet its precise functions remain unclear. Although dispensable for viral replication^34^, viruses lacking nsp2 showed slower vgRNA synthesis than WT viruses^33^. During early infection (DMVs ≲ 200 nm)^5^, SMLM in A549-ACE2 cells reveals nsp2 as ring-like shells adjacent to nsp3 that encapsulate dsRNA. This places nsp2 at DMV membranes and within convoluted membranes (CMs) between clustered DMVs, which is consistent with previous findings in MHV^5,7,8^ (Fig. 2C).

While the Golgi apparatus retains its typical interconnected ribbon morphology in early infection, it undergoes significant fragmentation in the late stage, consistent with previous reports of SARS-CoV-2–driven remodeling^15,35^ (Supplementary Fig. 2A, B). We confirmed that the Golgi matrix protein Giantin remains spatially distinct from ROs throughout this process (Supplementary Fig. 2A, B). Strikingly, however, nsp2 relocalizes to these fragmented Golgi structures (Fig. 2D). This specific Golgi association is, to our knowledge, novel and suggests evolving host-interaction roles for nsp2 during organelle remodeling.

### Nsp5 localizes to replication organelles rather than the bulk cytoplasm

Nsp5 cleaves the polyproteins pp1a/pp1ab at 11 sites between nsp4 and nsp16. Inhibition of the nsp5 protease prevents viral replication, making this protein an attractive drug target^36^. Among nsp5 inhibitors, nirmatrelvir (PF-07321332) is a clinically approved M^pro^ inhibitor for COVID-19 treatment^37^. Although many life-cycle schemes place polyprotein processing outside DMVs^16,38,39,40^, direct evidence has been limited.

In early infection, we observe a concentric localization of nsp5 around the dsRNA clusters, closer to dsRNA than nsp3 (Fig. 2E), suggesting a possible localization of nsp5 in the DMV lumen or on the DMV membranes. In late infection, the ROs/DMVs are better visible using vgRNA FISH staining that fills entire ROs (DMVs and vesicle packets)^5^. Our SR images show that nsp5 localizes in the same regions as vgRNA, indicating that nsp5 is associated with ROs in late infection, which is also illustrated by the bivariate pair-correlation functions (Fig. 2F). These observations suggest that, rather than functioning exclusively in the bulk cytoplasm as often depicted, the nsp5 protease might act within DMVs or on their surface.

### Radial/Angular super-resolution framework validates DMV pore nsp3–nsp4 geometry

DMV membrane pores contain the nsp3 and nsp4 proteins arranged in an approximately rotationally symmetric manner^41,42^. Nsp3 is located on the cytoplasmic side of the DMV, and its N-terminal domain extends ∼20 nm away from the outer DMV membrane^4^. Nsp4 is a much more compact protein, located close to the inner DMV membrane with its C-terminus facing the DMV interior. In this section, we aimed to precisely localize all SARS-CoV-2 NSPs within the DMV and with respect to the DMV membranes; therefore, we chose the position of the nsp4 C-terminus as a reference point for NSP localization. Because dsRNA fills the interior of DMVs^5,10^, we used the centroid of the dsRNA localizations as the origin point for the radial distance analysis of SARS-CoV-2 targets (Fig. 3A). We analyzed only cells in early infection stages with well-defined, isolated DMVs with a diameter of ≲200nm (Fig. 3B). In later infection stages, DMVs grow, merge into vesicle packets, and form less regular structures^5^, which would make this analysis more difficult.

**Fig. 3:**
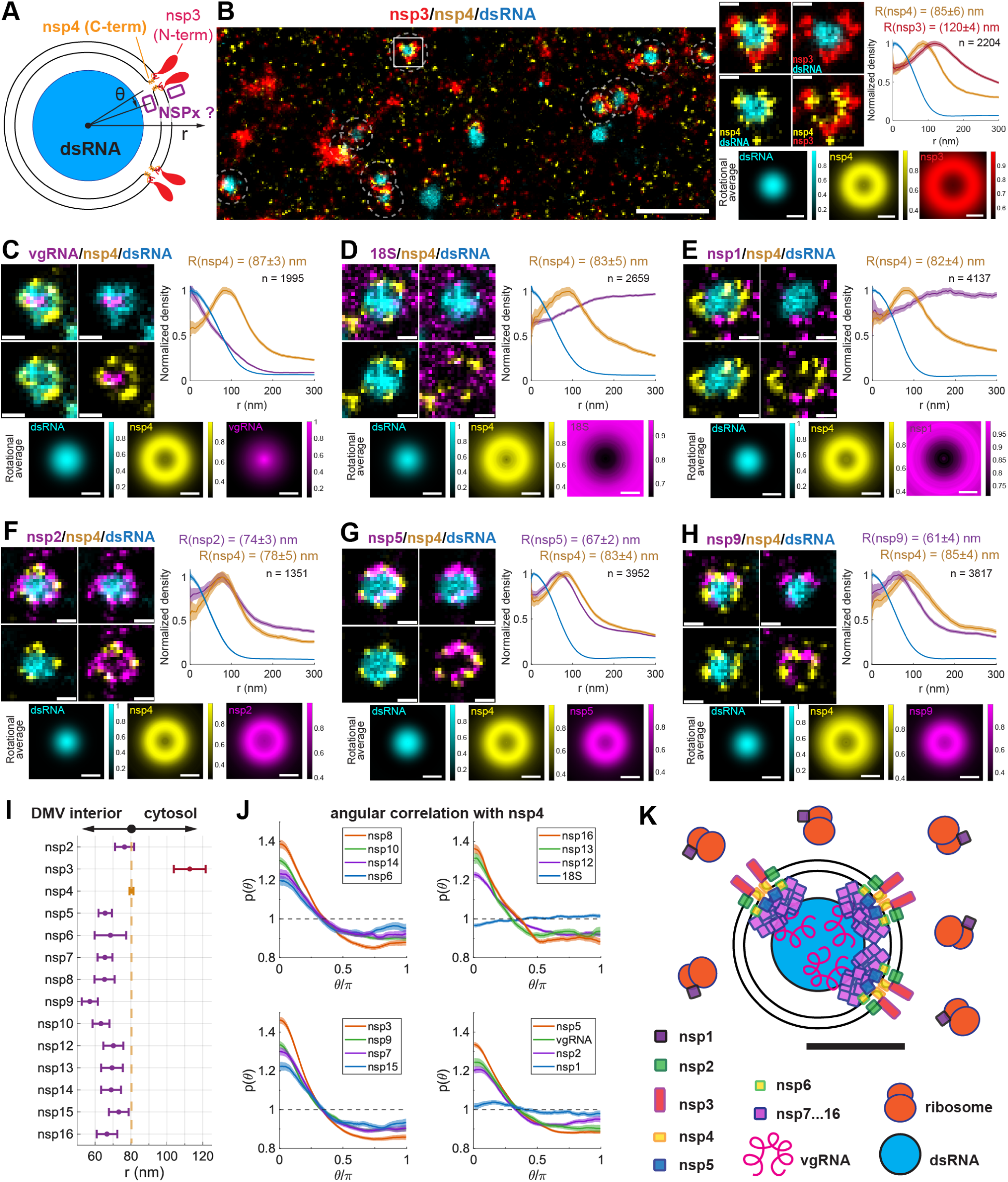
Nanoscale positioning of all SARS-CoV-2 NSPs with respect to DMVs. (A) Schematic of a SARS-CoV-2 DMV with dsRNA in the center and nsp4 with nsp3 at the membrane pores. The positions of all remaining components were determined with respect to the known positions of these reference targets. (B) Representative three-color SR image of the cytoplasm of a typical A549-ACE2 cell at an early infection stage. DMVs (dashed circles) appear as round dsRNA clusters (cyan) surrounded by nsp4 (yellow) and nsp3 (red). A magnified view of a single DMV (white box, top right) reveals the three-layer organization of dsRNA, nsp4, and nsp3. The corresponding radial density distribution functions, *g*(*r*) (top rightmost plot), and rotationally averaged images (bottom right) display the three-layer structure averaged across *n* DMVs from 16 different cells. (C–H) Representative three-color SR images of individual DMVs, corresponding radial density distribution functions, *g*(*r*), and rotationally averaged images for vgRNA, 18S rRNA, nsp1, nsp2, nsp5 and nsp9 relative to nsp4 and dsRNA. Data were obtained from 10 (C), 15 (D), 26 (E), 13 (F), 25 (G), and 22 (H) individual early-stage infected cells. Where applicable in (B–H), the average peak positions, *R*, of *g*(*r*), are indicated above the plots as the mean ± 95% CI, obtained via Gaussian fitting of the peak portion of *g*(*r*) following resampling with replacement (bootstrapping). *n* represents the total number of analyzed DMVs for each labeling combination. (I) Mean positions ± 95% CI for all NSPs associated with DMVs, calculated after rescaling the nsp4 positions within individual target groups (B–L) to match the global mean position of nsp4. The global position of nsp4 ± 95% CI (yellow error bar) was calculated across 13 groups (B, F–L) comprising n = 36,662 DMVs. Detailed data for the remaining NSPs are provided in Supplementary Fig. 4. (J) Angular bivariate pair-correlation functions, *p*(*θ*), of nsp4 with the indicated targets, represented as mean ± 95% CI. All targets peak at *θ* = 0, except 18S rRNA and nsp1. (K) Model of a mature, early-stage DMV based on the SR data. Scale bars: 1 µm (B, overview image), 100 nm (C–H, K and magnified individual DMVs and rotational averages in B).

The positioning of nsp3 and nsp4 is known precisely from the cryo-EM structure of the minimal DMV pore^4^. Therefore, to test our experimental approach, we first localized the N-terminal domain of nsp3 with respect to dsRNA and to the C-terminal domain of nsp4. The three-color SR images of dsRNA, nsp4, and nsp3 reveal a concentric three-layer structure: 1) a typical^5,10^ round dsRNA cluster in the center; 2) an nsp4 layer encapsulating dsRNA; and 3) a larger nsp3 layer that encapsulates both dsRNA and nsp4 (Fig. 3B). Next, for each DMV—defined as a dsRNA cluster surrounded by nsp4 localizations—we calculated a radial distribution function *g*(*r*) of the single-molecule localizations of various NSPs with respect to the centroid of the dsRNA cluster. Averaging over thousands of individual DMVs from multiple infected cells yielded a radial distribution, *g*(*r*), for each NSP with a distinct peak representing its average radial position within the DMV. Although the 2D projection of these 3D spherical structures introduces central density and a systematic inward shift of absolute peak positions, relative distances between concentric layers remain robustly preserved (Supplementary Fig. 3; see Methods). Relying on this relative measurement, our analysis reveals that the nsp3 N-terminus is positioned approximately 35 nm further from the DMV center than the nsp4 C-terminus (Fig. 3B), in excellent agreement with the known architecture of the DMV pore^4^.

Previously, it was reported that vgRNA localizes inside DMVs adjacent to dsRNA^5,7,10^. Indeed, we observe that within the DMV, vgRNA localizes on average at approximately the same radial positions as dsRNA (Fig. 3C), reflecting the accumulation of vgRNA in the interior of the DMV lumen^5^. Ribosomes are found only very rarely in DMVs^41,43^, suggesting that the translation of viral proteins mostly occurs outside ROs. Our analysis indicates that the average localization densities for both 40S ribosomal subunits and nsp1 are indeed lower in the ROs than in the surrounding cytosol (Fig. 3D, E), further confirming that DMVs are mostly not involved in viral protein synthesis.

### Nsp5 and replicase NSPs reside intraluminally and align with DMV pores for protease-dependent DMV maturation

The localization of SARS-CoV-2 nsp2 with respect to DMVs is largely unknown; however, there was evidence of nsp2 positioning at the cytosolic side of the DMV membrane of MHV^44^. The radial distribution function *g*(*r*) for nsp2 peaks at approximately the same radius as nsp4, indicating that SARS-CoV-2 nsp2 is also located very close to the membranes of early DMVs (Fig. 3F). Nsp6 is the third transmembrane protein among the coronaviral NSPs that transforms and zippers the host ER^45,46^ and therefore can be expected to anchor to DMV membranes. The *g*(*r*) for nsp6 shows that this protein is found slightly deeper into the DMV than nsp4 (Supplementary Fig. 4A), which suggests a possible nsp6 localization at the inner DMV membrane.

In related coronaviruses, nsp5 has long been shown to colocalize with nsp3^47^, to localize at convoluted membranes^8^, and to be protected by membranous structures^8,48^. However, the precise localization of nsp5 with respect to DMVs has not been previously measured. Our radial distribution analysis indicates that SARS-CoV-2 nsp5 localizes predominantly within the DMV lumen, close to the DMV membrane, on average 16 nm deeper than nsp4 (Fig. 3G). Because nsp5 is the main coronaviral protease, indispensable for viral polyprotein processing and for viral replication, its intraluminal localization provides direct spatial evidence for an emerging model of polyprotein processing coupled to DMV formation. We propose that the nsp5–nsp16 polyprotein block is co-enclosed during DMV biogenesis. In this scenario, partially uncleaved pp1a/pp1ab precursors are engulfed as the membrane seals, allowing nsp5 to subsequently complete the polyprotein processing within the protected DMV lumen.

The remaining nsp7–nsp16 proteins are involved in viral RNA replication and processing^39^ and therefore can be expected to localize and act within the protective DMV environment. Previous studies, including our earlier work^5^ and recent analogous super-resolution mapping in the alphacoronavirus HCoV-229E^10^, have localized the RdRp catalytic subunit nsp12 and the accessory domains nsp7, nsp8, and nsp9 to the DMV lumen. Furthermore, cryo-electron tomography (cryo-ET) studies have observed unassigned luminal densities positioned directly beneath the DMV pore^42^. However, high-resolution spatial data providing the molecular identities for the remaining non-structural proteins have been lacking. Using antibody labeling and SR microscopy, we measured the radial density distributions *g*(*r*) for nsp7/8/9/10/12/13/14/15/16 within DMVs. The peak in *g*(*r*) for these proteins lies approximately 5–25 nm deeper than for nsp4, suggesting that they are predominantly located in the DMV interior (Fig. 3H, Supplementary Fig. 4).

The average radial position of nsp4 exhibits some variability (73–88 nm) between different measurement pairs due to biological variability between the cells in different samples, which are inevitably found at slightly different infection time points. To precisely compare the positions of all NSPs, we aligned the average nsp4 position in each of the measurement pairs to the global average position of nsp4 calculated from all groups and applied the same scaling factor to the second target in each group. The resulting rescaled positions reveal that nsp5–nsp16 are found significantly deeper in the DMV than nsp4, while nsp2 localizes at a similar radial position to nsp4, and the N-terminus of nsp3 lies in the cytosol, significantly farther away from nsp4 (Fig. 3I).

Considering that nsp2–nsp16 lie near DMV pores radially, we asked whether they also align angularly around the RO boundary. We calculated the bivariate angular pair-correlation function^49^ *p*(*θ*) between nsp4 and each target (Fig. 3J). These functions reflect the average angular distance between two targets and would peak at *θ* = 0° if the two targets are systematically found at a similar angular position. *p*(*θ*) was normalized such that *p*(*θ*) = 1 for a non-correlated pair of species. Among all targets, *p*(*θ*) has the highest peak at *θ* = 0° for the nsp4 –nsp3 pair, reflecting the known colocalization of both targets in the DMV pore, which is directly noticeable in the SR images (Fig. 3B). 40S ribosomal subunits and nsp1 both demonstrate very weak angular correlation with nsp4, while vgRNA, nsp2, and nsp5–nsp16 strongly correlate with nsp4 in the angular direction, with a clear peak at *θ* = 0° (Fig. 3J). This indicates that nsp2 and nsp5–nsp16 predominantly localize close to the DMV pores not only in the radial but also in the angular direction around the membrane, suggesting that these proteins are attached to or located close to the pores within mature replication-competent DMVs, as indicated by the strong dsRNA signal in the core region.

To confirm the concentric localization of NSPs around the dsRNA core of DMVs/ROs and to rule out possible data misinterpretation due to 2D projection (three-color 2D dSTORM) of 3D objects (DMVs), we performed two-color 3D dSTORM microscopy using a double-helix point spread function (3D-DHPSF)^50^ on nsp5, nsp8, and nsp10 with dsRNA. This method allows for a near-isotropic resolution in 3D within a ∼2 µm imaging depth in the sample^19,51^. For each target pair, we indeed observe NSPs adjacent to dsRNA clusters and surrounding them in 3D (Supplementary Movie 1), which, when projected in 2D at different angles, resemble the ring-like 2D images of individual NSPs (Fig. 4A–C). The localization information derived from our experimental measurements is summarized in Fig. 3K.

**Fig. 4:**
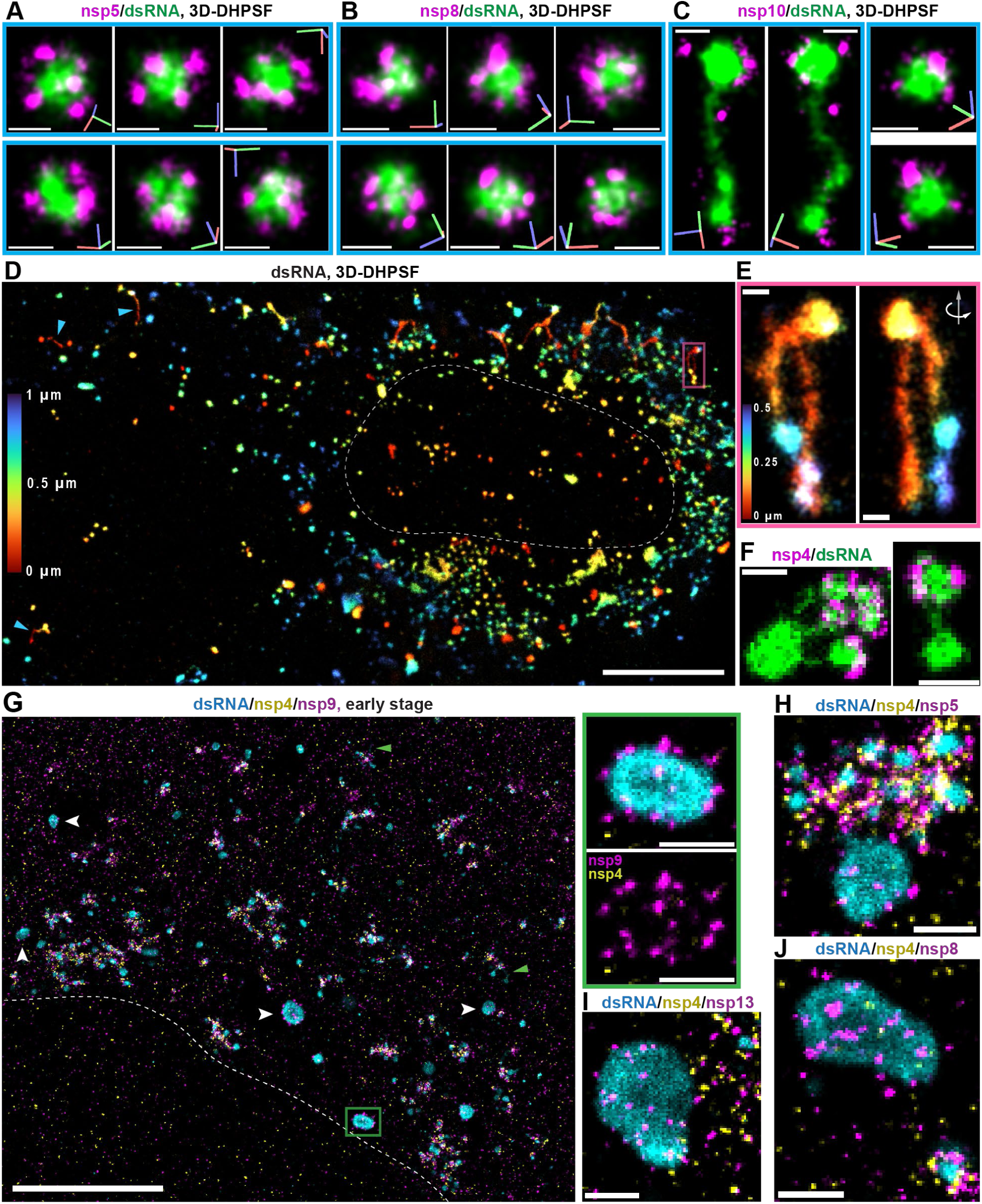
3D and 2D super-resolution architecture of DMVs and diverse dsRNA structures. (A–C) Representative 3D dSTORM images of DMVs acquired using a double-helix point spread function (DHPSF), colabeled for dsRNA and nsp5 (A), nsp8 (B), or nsp10 (C) in SARS-CoV-2–infected A549-ACE2 cells. Each blue-framed set displays the same DMVs from different viewing angles. Axes are indicated in the corner of each panel (red, X; green, Y; blue, Z). (D, E) 3D-DHPSF imaging of dsRNA in an infected cell. The magnified view in (E) corresponds to the pink box in (D). The axial (Z) position of the fluorophores is color-coded according to the provided scale, where 0 µm represents the coverslip. (F) 2D dSTORM images of dsRNA colabeled with nsp4 reveal thin dsRNA connectors between larger dsRNA structures. These images correspond to the yellow boxes in the whole-cell image shown in Supplementary Fig. 5A (G) Examples of a cell in an early infection stage that contains large, approximately round dsRNA granules decorated with nsp9 puncta but lacking an nsp4 shell. The magnified panels on the right correspond to the green box in the large image. Green arrowheads highlight additional examples of dsRNA connectors, and white arrowheads indicate selected dsRNA granules lacking nsp4. (H–J) Examples of large dsRNA granules largely lacking nsp4 but decorated with nsp5 (H), nsp13 (I), and nsp8 (J). White dashed curves denote the approximate nuclear edge. Scale bars: 5 µm (whole-cell images in D, G), 500 nm (H–J, and magnified panels in G), and 200 nm (A–C, E, F).

### dsRNA connectors route replication intermediates between DMVs and pore-free granules

We observe that some dsRNA clusters are interconnected by fine, fiber-like dsRNA strands (Fig. 4C, Supplementary Fig. 5A) which we term “connectors”. The width of the dsRNA connectors, measured as *σ* of an equivalent Gaussian function, is only (17 ± 5) nm (Supplementary Fig. 5A), meaning it is limited by our single-molecule localization precision (*σ*_loc_ ≈ 15 nm, quantified from ThunderSTORM fits) and by the size of the antibodies used for dsRNA detection (up to ∼20 nm, primary + secondary antibodies^52^). This suggests that the dsRNA “connectors” might potentially be helical dsRNA molecules residing in the lumen of ER tubules, as such tubules commonly interconnect DMVs as observed by EM studies^42^. In addition to human A549-ACE2 cells, we also observed these dsRNA fibers in the Vero E6 cell line (Supplementary Fig. 5B), indicating that these fibers are a general feature of intracellular SARS-CoV-2 infection. These dsRNA connectors are also easily visible in 3D SR microscopy (Fig. 4C–E, Supplementary Movie 2).

In the cytoplasm of SARS-CoV-2–infected cells, alongside typical dsRNA clusters (d ≲ 200 nm) surrounded by nsp3, nsp4, and other NSPs (Fig. 2, Fig. 3), we also observe larger dsRNA clusters (d ≳ 250 nm) that lack the DMV membrane pore proteins nsp3 and nsp4 (Fig. 4G–J). These granules or condensed structures typically remain approximately round; however, in cells at later infection stages they may adopt less regular shapes and may occupy larger areas of the cytoplasm (Supplementary Fig. 5C–G). Despite the absence of nsp3 and nsp4, these aggregates are typically decorated with sparse puncta of various NSPs, including nsp9 (Fig. 4G, Supplementary Fig. 5C), nsp5 (Fig. 4H, Supplementary Fig. 5D), nsp13 (Fig. 4I), nsp8 (Fig. 4J), nsp10 (Supplementary Fig. 5E) and nsp12 (Supplementary Fig. 5F). However, these dsRNA structures mostly lack nsp2 (Supplementary Fig. 5D) and are located in the cytoplasmic regions devoid of nsp1 (Supplementary Fig. 5G). The existence of dsRNA aggregates with attached replicase proteins but lacking nsp3 and nsp4 indicates that the replicase proteins might possibly exercise their functions also without interaction with DMV pores. In some cases, larger nsp3/nsp4-lacking dsRNA structures are linked to several normal DMVs by dsRNA “connectors” (Fig. 4F, Supplementary Fig. 5A), suggesting a hypothetical mechanism of RO filling and viral RNA propagation in the cytoplasm via dsRNA “runners”.

### M protein maps the secretory–lysosomal route of SARS-CoV-2 assembly

Following the synthesis and spatial routing of vgRNA at the replication organelles, the viral life cycle shifts to packaging and egress. While this phase is governed by four main structural proteins—spike (S), membrane (M), envelope (E), and nucleocapsid (N)^53,54^—we focused our investigation on the nanoscale localization of the S, M, and N proteins to complete our spatial atlas. First, we validated the antibodies using virion labeling and SR imaging, which revealed a concentric localization of the S and M proteins, with the N protein located mostly in the center of the virion (Supplementary Fig. 6A). The diameter of the S ring in the labeled virions was 86 nm, while for the M protein this diameter was 18 nm smaller, consistent with the virion dimensions and with the positioning of M in the viral envelope and S on the exterior^55^.

M is the most abundant structural protein in the SARS-CoV-2 virion membrane and plays critical roles in viral particle assembly and egress^54^. In A549-ACE2 cells at 24 hpi, M protein antibody staining and SR imaging revealed four morphological structures exhibiting distinct localization patterns from the perinuclear region to the cytoplasmic membrane, possibly corresponding to different stages of viral assembly and egress (Fig. 5A, B): (i) perinuclear Golgi-proximal M proteins show large unstructured conglomerates where the M distribution is relatively sparse (Fig. 5A, blue square); (ii) mid-cytoplasm M proteins show small, dense aggregates (Fig. 5A, green square); (iii) peripheral cytoplasm M proteins form virion-sized particles that aggregate into large hollow clusters (Fig. 5A, red square); (iv) plasma membrane M proteins form virion-sized particles with M distributed on the viral envelope (Fig. 5B).

**Fig. 5:**
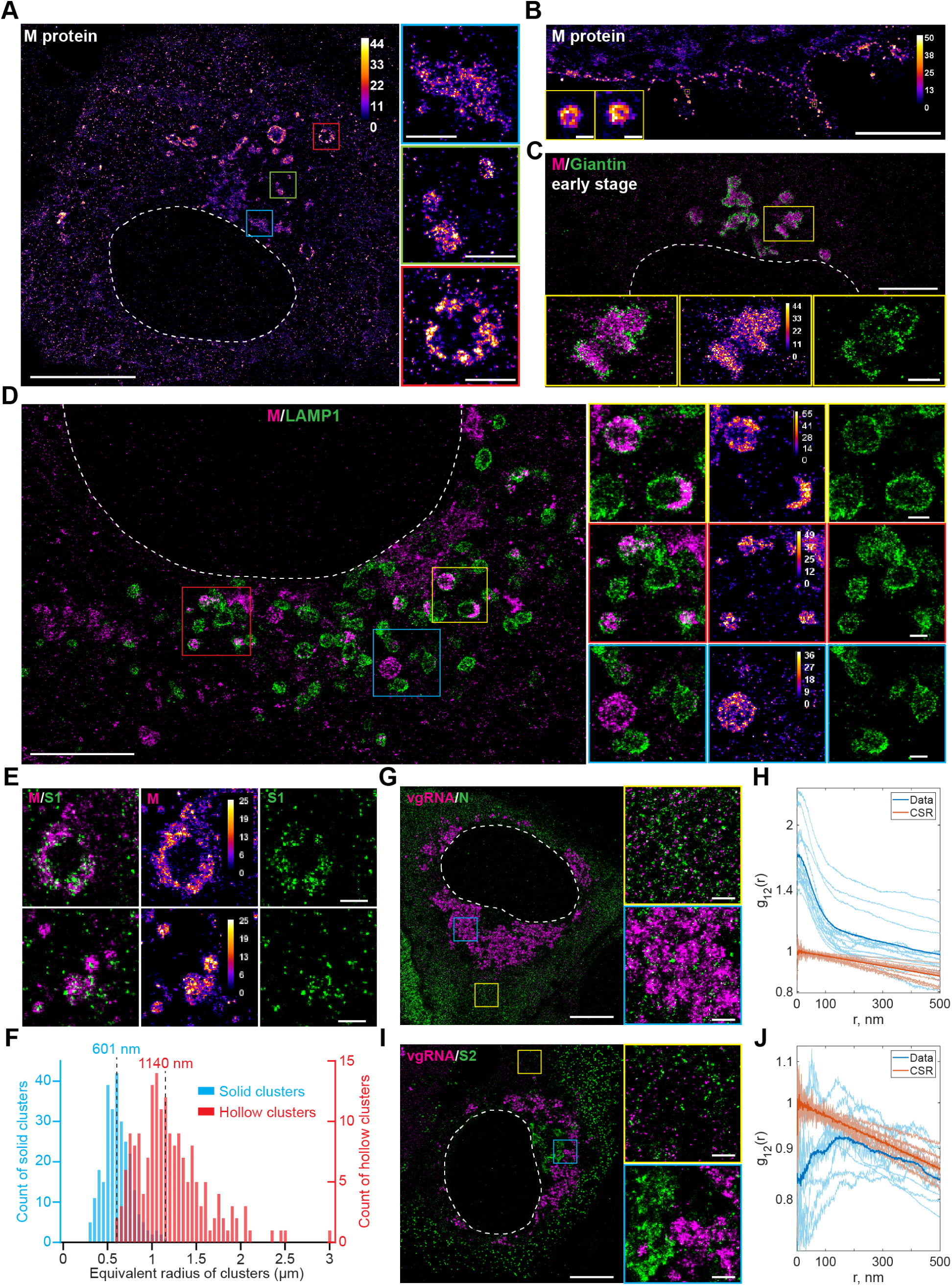
Nanoscale localization of the SARS-CoV-2 structural proteins M, S, and N. (A) Representative SR image of the M protein reveals distinct morphological structures. Magnified panels on the right correspond to the blue, green, and red boxes in the main image. (B) Representative SR image of the M protein shows virions arrayed on the cell membrane. Insets (yellow boxes) display magnified individual virions. (C) Representative two-color SR image of M protein (magenta) and the Golgi marker Giantin (green) indicates that M protein localizes within the Golgi apparatus. Bottom panels show magnified views of the region enclosed by the yellow box. (D) Representative two-color SR image of M protein (magenta) and the lysosomal marker LAMP1 (green) details the distribution of M protein inside and outside lysosomes. Panels on the right show magnified views of the regions enclosed by the corresponding colored boxes. (E) Representative two-color SR images demonstrate that M protein (magenta) codistributes with S1 protein (green) in both hollow (top) and solid (bottom) clusters. (F) Histogram of the equivalent radius of solid (blue, n = 276 clusters) and hollow (red, n = 180 clusters) M protein structures reveals two distinct size populations. Data were obtained from 22 cells and 8 independent experiments. (G) Representative two-color SR image of vgRNA (magenta) and N protein (green) at 24 hpi demonstrates colocalization exclusively in the cytoplasm outside ROs. Panels on the right show magnified views of the regions enclosed by the yellow and blue boxes. (H) Bivariate pair-correlation functions, *g_12_*(*r*), calculated between vgRNA and N protein localizations in the cytoplasm outside ROs indicate their close spatial association. (I) Representative two-color SR image of vgRNA (magenta) and S2 protein (green) shows a lack of colocalization both within and outside ROs. Panels on the right show magnified views of the regions enclosed by the corresponding colored boxes. (J) Bivariate pair-correlation functions, *g_12_*(*r*), calculated between vgRNA and S2 protein localizations in the cytoplasm outside ROs indicate a nanoscale spatial anti-correlation between them. White dashed lines in A, C, D, G, and I denote the approximate edge of the cell nucleus (large dark region). Color bars in A, B, D, E indicate the number of single-molecule localizations per SR pixel (20 × 20 nm^2^, except for B, 16 × 16 nm^2^). Scale bars: 10 µm (large image in A), 5 µm (large images in B–D, G, I), 1 µm (magnified images in A, C), and 500 nm (E and magnified images in D, G, I). Images and data in A–F were obtained from SARS-CoV-2–infected A549-ACE2 cells, whereas those in G–J were obtained from infected Vero E6 cells. All cells were fixed at 24 hpi.

To explore these diverse morphological structures of the M protein, we co-imaged M with host compartments. Since M is glycosylated^56^ and most glycosylation occurs in the Golgi, we costained M with an anti-Giantin antibody that labels the rim of Golgi cisternae^57^. We found that the large, unstructured conglomerates of M protein distributed in the perinuclear region localize within the Golgi (Fig. 5C), which is likely essential for M glycosylation. Super-resolution imaging reveals that the cis/medial-Golgi marker Giantin forms a peripheral rim around these M clusters, suggesting accumulation within the cisternae. To map the exit route, we co-imaged the trans-Golgi network (TGN) marker TGN46. Unlike the encapsulation seen with Giantin, M aggregates appear closely juxtaposed to or partially overlapping with TGN46-positive structures (Supplementary Fig. 7A), consistent with the protein traversing the TGN for sorting and packaging.

We then costained M with an anti-LAMP1 antibody that labels the membrane of lysosomes and late endosomes^58^, as it has been reported that β-coronaviruses—including SARS-CoV-2—use lysosomes for egress^58^. Indeed, some lysosomes contained dense M aggregates, while many lacked M (Fig. 5D). Interestingly, in large lysosomes, M lined the inner lysosomal membrane, producing a hollow appearance (Fig. 5D, yellow); in smaller lysosomes, M filled the lumen (Fig. 5D, red). We also observed large M clusters outside LAMP1 compartments (Fig. 5D, blue). To assess whether these M-positive aggregates correspond to assembled virus particles, we colabeled the M protein alongside the spike S1 subunit. This revealed the co-assembly of both proteins into virion-sized particles that pack into large, hollow clusters (Fig. 5E). Our findings suggest that these nanoscale particles represent either fully assembled virions or late-stage intermediates undergoing assembly. However, we cannot exclude the alternative possibility that they correspond to mature virions re-internalized into lysosomes via endocytosis. We further quantified the size of different M clusters and found that small, dense M clusters have a median equivalent radius of 601 nm and large, hollow M clusters have a median equivalent radius of 1140 nm (Fig. 5F), comparable to lysosome dimensions (0.5–1 µm)^59^.

### Spike subunits segregate after furin cleavage and N–vgRNA RNPs form outside DMVs

Next, we explored the nanoscale cellular localization of the N proteins and S proteins by co-imaging with vgRNA. In early infection, N proteins tend to accumulate in the cytosol outside and around ROs (Supplementary Fig. 6C). This observation is consistent with previous reports^5,9,60,61^ and is potentially reflecting the encapsulation of newly synthesized vgRNA as it exits the DMV pore. In late infection, N proteins distribute more evenly in the cytoplasm (Fig. 5G) as vgRNA moves to assembly sites and disperses into the cytoplasm. We observed that a subset of cytoplasmic N proteins colocalized with vgRNA puncta, while a fraction remained unassociated (Fig. 5G, yellow square, and 5H). This suggests that N proteins not only assemble with vgRNA into ribonucleoprotein particles (RNPs), but may also establish a pool that performs distinct functions in the cytoplasm^62^.

S proteins detected via the S2 domain antibody do not localize inside ROs, and their distribution in the cytoplasm is heterogeneous (Fig. 5I, Supplementary Fig. 6B). Namely, S2 proteins aggregate to form clusters within the cytoplasm that do not colocalize with vgRNA puncta (Fig. 5I–J). Similar to M, perinuclear S2 localizes to the Golgi starting in early infection (Supplementary Figs. 6B, 7B). Consistent with M protein trafficking, S2 clusters are encapsulated by Giantin but localize adjacent to TGN46-positive structures (Supplementary Fig. 7B), indicating coordinated transit through the secretory pathway.

We also detect S2 subunits in ER tubules (labeled by Sec61β-GFP overexpression) and decorating the nuclear envelope (Supplementary Fig. 6D; antibody specificity confirmed in Supplementary Fig. 8). Because host cell furin-like proteases likely cleave the spike protein^63^, we simultaneously colabeled the S1 and S2 domains. While both domains localize to the Golgi and to vesicles with virions (Fig. 5E, Supplementary Fig. 6E), S1—unlike S2—does not localize to ER tubules or the nuclear envelope (Supplementary Fig. 6E). This suggests that the SARS-CoV-2 S protein partially disassembles after the furin cleavage in the host cells, with a fraction of S2 domains remaining in various cellular membranes while S1 domains follow the standard virion assembly pathway to form surface-anchored spikes.

### Protease inhibition by nirmatrelvir generates multilayer polyprotein bodies that persist and precede rebound

After defining the baseline infection landscape in A549-ACE2 cells, we perturbed the viral life cycle with the nsp5 (M^pro^) inhibitor nirmatrelvir (PF-07321332), the active component of Paxlovid^37^. Preventing the polyprotein cleavage at all M^pro^ sites leaves pp1a/pp1ab uncleaved between nsp4 and nsp11/nsp16; only nsp1, nsp2 and nsp3 are released via PL^pro^ (within nsp3) cleavage (Fig. 1C). Given that nsp7–nsp16 are required for viral RNA replication and transcription, inhibiting M^pro^ should halt viral replication. However, it remains unknown how this treatment affects the transformation of host ER into DMVs, as this process can be executed solely by nsp3 with nsp4^41^. Inhibiting M^pro^ cleavage with nirmatrelvir also provides a tool to investigate DMV maturation by trapping the process in an intermediate state with uncleaved polyproteins.

We treated infected cells (MOI = 2) with three concentrations of nirmatrelvir (0.1×, 1×, and 10× EC_50_; EC_50_ = 144 nM) at 0, 2, or 6 hpi, and quantified the percentage of infected cells at 24 hpi using confocal imaging of immunofluorescence staining of dsRNA, nsp4, and nsp5 (Supplementary Fig. 9A). Treatment with 14.4 nM nirmatrelvir yielded no decrease in infected cells at any time points. Conversely, treatment cells with 1440 nM at 0 and 2 hpi nearly abolished infection. Treatment with 1440 nM at 6 hpi, or 144 nM at any time point, produced a significant (>2.6-fold) reduction in infected cells compared to the untreated control (Supplementary Fig. 9A).

While cells treated with 14.4 nM nirmatrelvir exhibited no noticeable phenotype change in DL or SR, approximately 33% (11 out of 33 imaged cells, compared to 0 out of 7 in the untreated group) of SR-imaged infected cells treated with 144 nM nirmatrelvir at any time point, or with 1440 nM at 6 hpi, displayed multilayered bodies (MLBs) in the cytoplasm (Fig. 6, Supplementary Fig. 9B–D). When labeled for nsp4 and nsp5 and imaged using 2D dSTORM, these MLBs exhibit ring-like or crescent-like shapes with dimensions of 0.5–5 µm (Fig. 6A, Supplementary Fig. 9B). These bodies lack a significant dsRNA signal, indicating they are not associated with active vgRNA replication. Within the MLBs, we resolved several layers containing both nsp5 and nsp4, with a spacing of ∼75 nm between layers (Fig. 6A, Supplementary Fig. 9B). We then labeled the C-terminus of pp1a (nsp10) and C-terminus of pp1ab (nsp16) and found that both proteins also locate to the MLBs (Fig. 6B–C). This indicates that these layers likely contain the uncleaved pp1a/pp1ab polyproteins (comprising nsp4–nsp10/nsp16) resulting from M^pro^ inhibition by nirmatrelvir.

**Fig. 6:**
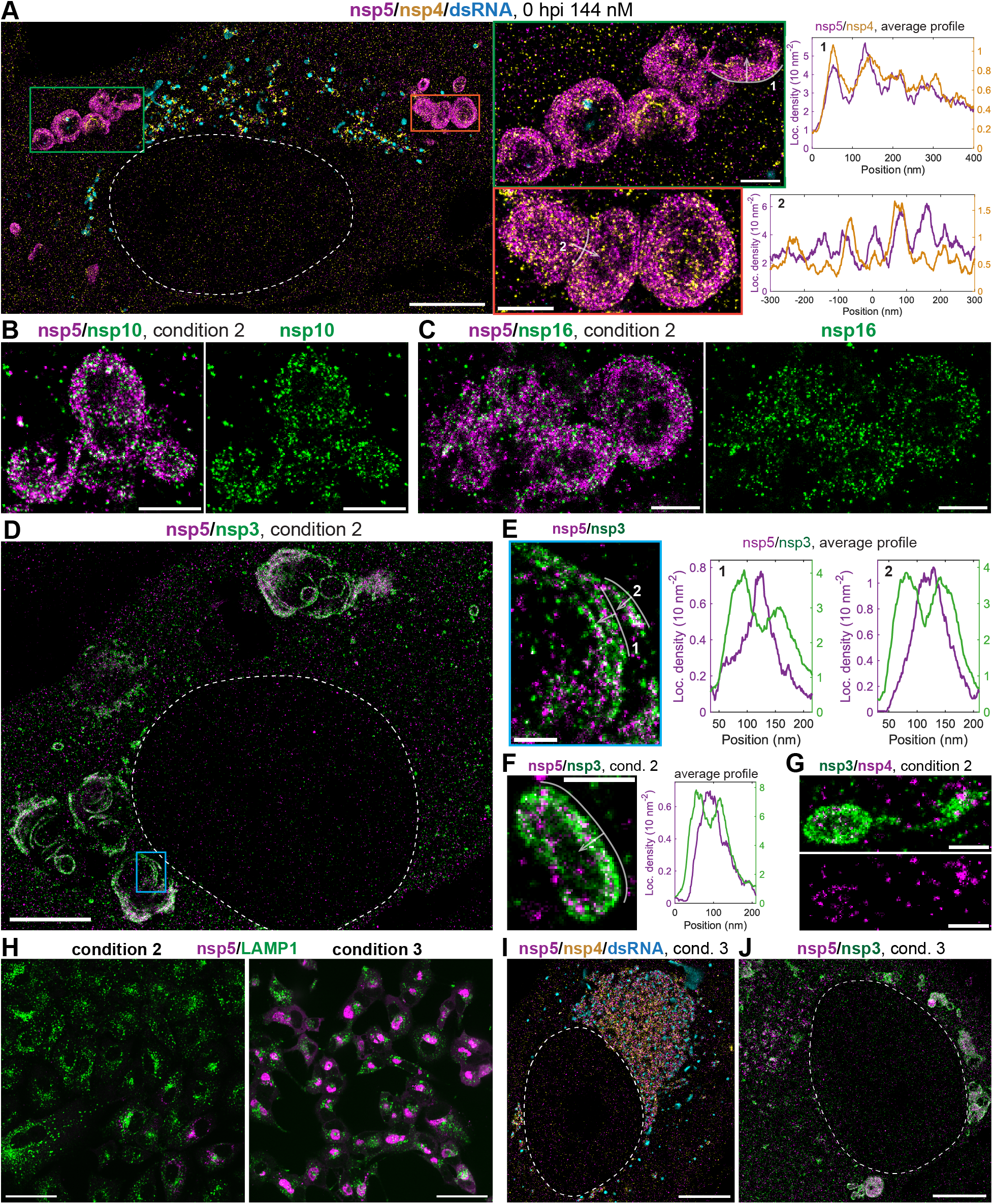
Multilayered bodies in SARS-CoV-2–infected cells treated with nirmatrelvir. (A) Representative SR image of an infected cell (MOI = 2) treated with 144 nM nirmatrelvir from 0 to 24 hpi and fixed at 24 hpi. Two regions with large multilayered bodies (MLBs) are magnified in the central panels. Density profiles (right, panels 1 and 2) show the localization density of nsp4 and nsp5 calculated perpendicular to the corresponding numbered white curves, in the direction indicated by the arrows (localization density orthogonal to the arrows was averaged). Both nsp4 and nsp5 localize in layered structures with a periodicity of ∼75 nm. (B–G) SR images of infected cells (MOI = 10) treated with 288 nM nirmatrelvir from 6 to 24 hpi and fixed at 24 hpi. Nsp10 (B), nsp16 (C) and nsp3 (D) localize to the same multilayered structures as nsp5. (E) Magnified view of the region in (D) demonstrates the localization of nsp3 on both sides of nsp5-labeled layers. The averaged density profiles of nsp3 and nsp5 (right), calculated perpendicular to the numbered white curves in the direction of the arrows, reveal a double structure of nsp3 layers centered on single nsp5 layers. (F,G) Additional examples of MLBs from different cells in the same sample group, exhibiting this double nsp3 layers structure. (H) Confocal images of infected cells (MOI = 10) treated with 288 nM nirmatrelvir from 6 to 24 hpi and fixed at either 24 hpi (left) or 48 hpi (right), utilizing lysosome-associated membrane protein 1 (LAMP1, green) as a general cellular marker. In the left panel, only 6 cells in the bottom right corner display a weak punctate nsp5 signal (magenta) typical for an early infection phenotype, whereas the remaining cells show only LAMP1 signal and no detectable nsp5. In the right panel, nearly all cells exhibit a strong perinuclear nsp5 signal consistent with late-stage infection. (I, J) SR images of infected cells (MOI = 10) treated with 288 nM nirmatrelvir from 6 to 24 hpi and fixed at 48 hpi. A typical cell under these conditions displays a late-infection phenotype with highly dense perinuclear NSP and dsRNA localizations (I), whereas another typical cell shows prominent cytoplasmic MLBs (J). Among 32 infected cells imaged in SR at 48 hpi, 16 (50%) contained MLBs. In contrast, among 41 similarly treated infected cells fixed at 24 hpi (examples shown in B–G), 29 (∼71%) contained MLBs. White dashed curves denote the approximate nuclear edge. Scale bars: 50 µm (H), 5 µm (large images in A; D, I, J), 1 µm (middle panels in A; B, C), and 500 nm (E–G). All images and data were obtained from SARS-CoV-2–infected A549-ACE2 cells.

To investigate whether MLBs result from altered DMV biogenesis, we colabeled the essential pore proteins nsp3 and nsp4. We found that nsp3 localizes to MLBs and forms a double-layered structure, where the N-terminus of nsp3 is positioned ∼30 nm away from each nsp4/nsp5 layer on both sides (Fig. 6D–G, Supplementary Fig. 9C–D). This offset of the nsp3 N-terminus from nsp4/nsp5 matches the distance of the cytoplasmic nsp3 N-terminus from standard DMV membranes (Fig. 3B). This double-layered pattern suggests that—unlike in canonical DMVs where nsp3 is restricted to the cytoplasmic side of the double membrane—nsp3 in nirmatrelvir-induced MLBs sits on both sides of zippered, locally planar membranes.

Alongside MLBs, we also observed morphologically normal DMVs in the nirmatrelvir-treated cells (Fig. 6A, Supplementary Fig. 9B–E). Containing dsRNA and NSPs, these remaining DMVs are likely replication-competent, providing the vgRNA necessary for pp1a/pp1ab translation and for the further formation of multilayered aggregates. This observation provides a mechanism for viral rebound, consistent with the clinical recrudescence observed after Paxlovid treatment.

To test this, following nirmatrelvir incubation from 6 hpi to 24 hpi, we performed a drug washout and cultured the cells for an additional 24 h without adding new virus. While few cells exhibited viral target signals at 24 hpi, nearly all cells displayed robust infection by 48 hpi (Fig. 6H, Supplementary Fig. 9A). These cells displayed a predominant late-infection phenotype with intense perinuclear NSP and dsRNA signals (Fig. 6H). SR imaging revealed canonical late-infection morphologies with dsRNA clusters and very dense NSPs in the perinuclear region (Fig. 6I), suggesting that vgRNA persisted during nirmatrelvir inhibition, enabling the virus to rapidly resume replication once the M^pro^ inhibitor was removed. Many cells still contained MLBs (Fig. 6J), meaning that the cell machinery cannot fully degrade these structures during the 24 h post-washout period.

Together, these data indicate that M^pro^ blockade arrests DMV maturation and drives the formation of multilayered, NSP-decorated polyprotein assemblies. However, since a subset of DMVs remains functional, rapid viral rebound occurs once inhibition is lifted. Therapeutically, achieving complete suppression may require eliminating residual ROs and vgRNA, or destabilizing MLBs to prevent recrudescence.

Considering that preventing cleavage of the nsp3–nsp4 junction could also lead to a comparable phenotype of limited DMV formation observed under nirmatrelvir treatment, we next performed Western blot studies to determine whether nsp3–nsp4 cleavage occurs under nirmatrelvir, which is critical to validating our models of DMV and MLB formation. We probed nsp3 using an anti-nsp3 antibody in Western blot analysis of lysates from SARS-CoV-2-infected cells treated with nirmatrelvir or DMSO. Specific detection of nsp3 was confirmed, by the absence of its signal in the uninfected control group (Supplementary Fig. 10A–B, Virus−). The expression level of nsp3 was significantly decreased after nirmatrelvir treatment (Supplementary Fig. 10C), while the proportion of uncleaved nsp3 (i.e., nsp3+) did not significantly change between DMSO and nirmatrelvir treated groups (Supplementary Fig. 10D), indicating that nirmatrelvir treatment has no significant influence on the cleavage between nsp3 and other proteins, including nsp4. Since the nsp3–nsp4 junction is cleaved by the viral papain-like protease (PL^pro^, located within nsp3) rather than M^pro^, this confirms that nsp3 is released from nsp4 even when nsp5 is fully inhibited by nirmatrelvir.

## Discussion

Despite intensive study, key cellular steps of coronavirus infection remain unresolved and may underlie the wide range of clinical severity. Using super-resolution mapping in human cells, we localize key SARS-CoV-2 components (Fig. 7A; Supplementary Table 2) with nanometer precision to test and refine models of replication, organelle remodeling, and assembly.

**Fig. 7:**
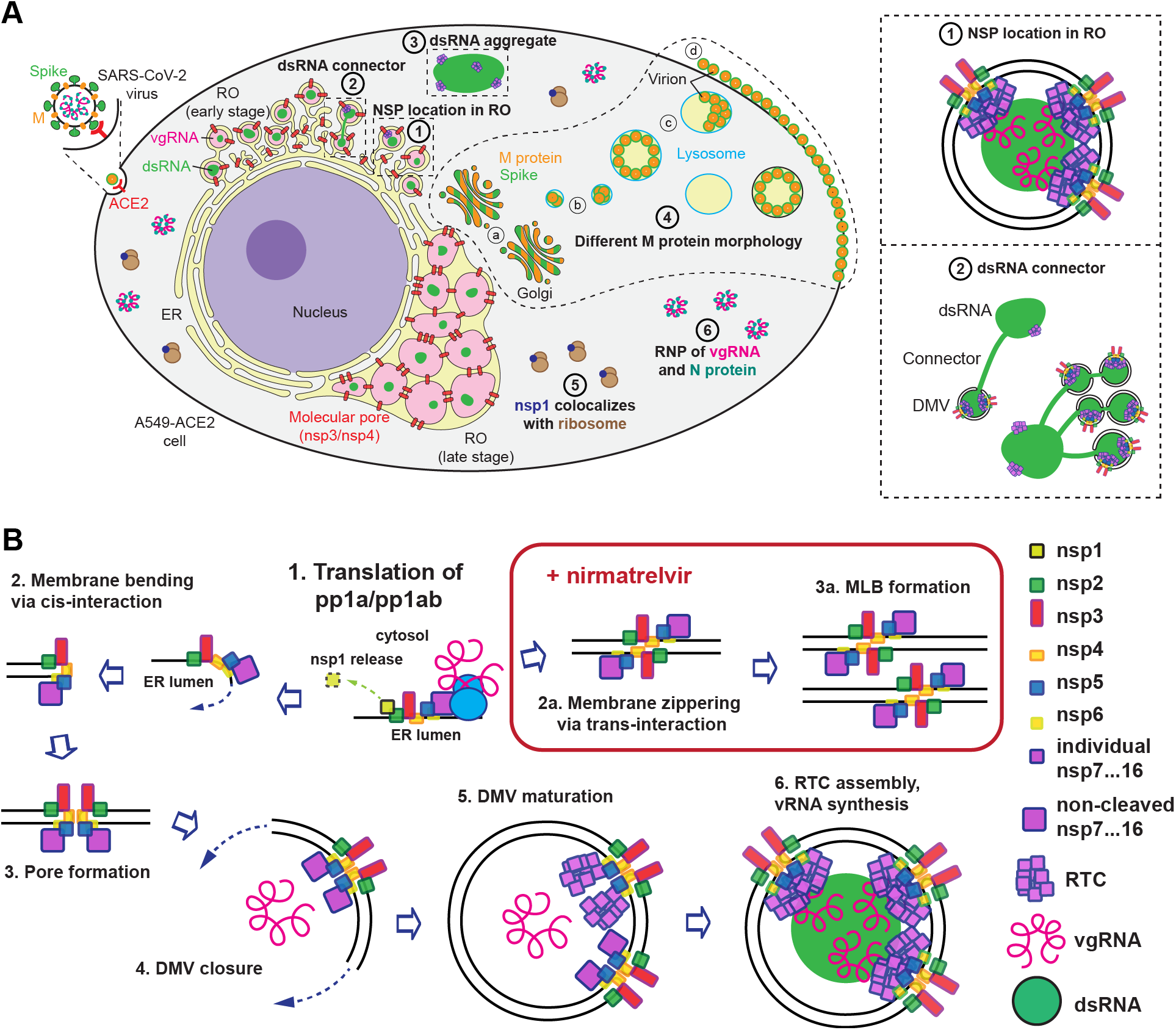
Summary of findings from the super-resolution spatial Atlas of SARS-CoV-2 infection. (A) Schematic summarizing the novel viral infection landscapes revealed from the SR Atlas in human cells. (1) Nanoscale localization of NSPs within ROs is detailed in the top right inset. (2) Fiber-like connectors physically link distinct dsRNA clusters, as detailed in the bottom right inset. (3) Large, nsp3/nsp4-lacking dsRNA aggregates (granules) are decorated with replicase components. (4) M protein exhibits diverse morphological structures across various cellular regions: (a) colocalizing with S protein within the Golgi; (b) forming small solid clusters (putative virions assembled with spike) inside lysosomes; (c) forming large hollow clusters inside lysosomes and other vesicles (putative virions assembled with spike); (d) assembled into virions arrayed on the plasma membrane. (5) Nsp1 colocalizes with ribosomes in the cytoplasm. (6) vgRNA colocalizes with N protein to form ribonucleoproteins (RNPs) in the cytoplasm. (B) Proposed model of DMV formation, polyprotein processing, and disruption by nirmatrelvir: (1) The process begins with vgRNA translation into polyproteins pp1a/pp1ab on the endoplasmic reticulum (ER) membrane. The polyprotein is anchored by the transmembrane domains of nsp3, nsp4 and nsp6. Nsp1 is cleaved by the PL^pro^ domain of nsp3 shortly after translation, while M^pro^ (nsp5) possibly auto-cleaves and begins processing the C-terminal domains (nsp5–11/16). (2) This early cleavage allows the *cis*-interaction of the nsp3 and nsp4 ectodomains, leading to membrane bending, while nsp5–11/16 stays attached to the nsp4 C-terminus via non-covalent and remaining covalent bonds. (3) This is followed by pore formation^4^, which partitions nsp2 and nsp3 to the cytosolic face of the pore, while the maturing nsp5–11/16 proteins are compartmentalized on the opposite side of the growing double membrane. (4) The DMV membrane closes, encapsulating vgRNA; the binding of vgRNA to the nsp7–nsp11/16 complexes ensures vgRNA localization within the DMV interior. (5) Within the closed DMV, optimal local concentrations of nsp5 complete the cleavage of pp1a/pp1ab to produce a mature RO. (6) Replication-transcription complexes (RTCs) assemble from NSPs to replicate, transcribe and modify viral RNAs within the DMV. (2a) The addition of nirmatrelvir inhibits M^pro^-mediated polyprotein cleavage. The retention of the completely uncleaved nsp4–11/16 polyproteins likely prevents proper membrane bending and instead promotes *trans*-interactions between nsp3 and nsp4 ectodomains. This results in the zippering of adjacent ER membranes, which eventually assemble into aberrant multilayered bodies (MLBs) (3a).

The absence of ribosomes in the DMV interior (Fig. 3D) indicates that protein translation occurs in the cytoplasm, not within ROs. While schematic models routinely depict the processing of pp1a/pp1ab by PL^pro^ and 3CL^pro^ occurring entirely in the cytoplasm^16,38,39,40^, much evidence places vgRNA replication and the mature RTCs inside the DMVs^5,6,7,10^. Because the central channel of the DMV pore (width < 20 Å^4^) is too narrow for large, folded proteins to traverse, it has recently been hypothesized that the large coronaviral NSPs must be enclosed at the time of DMV formation^9,10^. Our finding that nsp5 (3CL^pro^) localizes intraluminally, near the inner membrane and pores in early DMVs (Fig. 2E, F, 3G, 3I), provides direct experimental validation for this emerging model.

We propose that polyprotein processing is dynamically coupled to DMV biogenesis. In this model, nsp3–nsp4 interactions initiate membrane bending while nsp4 remains spatially coupled to the partially uncleaved nsp5–nsp11/16 block within pp1a/pp1ab (Fig. 7B). This coupling during DMV biogenesisis is likely essential for luminal delivery, as it has been shown that when mature replicase NSPs are expressed individually—outside the context of the polyprotein—they completely fail to translocate into pre-formed DMVs^12^. Recent computational modeling independently supports a similar sequence in which pore formation precedes nsp5-mediated cleavage and traps the replicase inside the DMV^11^. Once initial cleavages relieve steric constraints to permit membrane closure, a subsequent trigger (e.g. the increased concentration of NSPs, which might favor the nsp5 dimerization necessary to increase its activity^64^) would stimulate 3CL^pro^ to finish DMV maturation. Analogous cleavage-gated switches operate in other viruses (e.g., HIV Gag maturation)^65^.

High-resolution structural data about the DMV pore are available only for the minimal nsp3–nsp4 pore without infection, and cryo-EM averaging would obscure flexible regions such as nsp5–nsp11/16^4^. Nevertheless, the C-terminal domain of nsp4, where nsp5 would be attached in the uncleaved polyprotein, is oriented toward the DMV interior^4^, which is compatible with our model of polyprotein cleavage after DMV formation. The N-terminal domain of nsp3 is located in the cytoplasm^4^; therefore, the PL^pro^ cleavage of nsp1 and nsp2 from the nsp3 N-terminus would release them directly into the cytosol. Nsp1 indeed has a cytoplasmic localization (Fig. 3E), and a significant part of nsp2 also localizes outside DMVs, in the CMs and the Golgi (Fig. 2C, D). DMV-associated nsp2 localizes very close to the membrane (Fig. 3F, I). By analogy with MHV^44^ and predicted transmembrane domains^66^, it likely occupies the cytosolic face of the outer DMV membrane rather than the lumen, consistent with its PL^pro^ cleavage in the cytosol and Golgi relocalization (Fig. 2D).

Together, our data support a protease-dependent maturation model in which nsp5–nsp16 accumulate inside DMVs, making the DMV lumen a protected chamber for viral RNA replication, transcription, and storage. By leveraging the high spatial resolution of dSTORM (∼15–20 nm), our study expands upon recent STED/SIM observations in the alphacoronavirus HCoV-229E^10^ to encompass the entire SARS-CoV-2 replicase. Furthermore, by incorporating not only radial but also angular pair-correlation analysis with respect to the pore complex, our data provide direct molecular confirmation for the unassigned RTC densities observed beneath the pore in cryo-ET data^42^, offering a comprehensive, detailed map of the mature replication organelle.

Recognizing that SMLM yields static snapshots of an asynchronous infection, the diverse compositional states we observe reflect an ongoing organelle maturation trajectory. As we previously established using time-controlled infections^5^, DMV maturation proceeds through distinct phases defined by RNA synthesis. Following the M^pro^-gated membrane closure described here, early-stage DMVs undergo rapid negative-strand synthesis, forming the dense dsRNA cores visualized in our structural atlas. Given that dsRNA accumulation saturates early in infection, the transition to late-stage DMV progression is instead characterized by the massive production and intraluminal accumulation of positive-sense vgRNA, which drives subsequent DMV expansion^5^. The canonical, pore-containing DMVs captured in our current high-resolution datasets predominantly represent this early-to-intermediate active replication phase, prior to extensive vgRNA-induced swelling.

Alongside these canonical structures, our atlas reveals a secondary, compositionally distinct class of dsRNA compartments that appear to form outside the primary maturation pathway. We observed massive, spherical dsRNA granules—sometimes exceeding 1 µm in diameter—that completely lack the DMV pore proteins (nsp3/nsp4) and cytoplasmic modulators (nsp1/nsp2) (Fig. 4F–J, Supplementary Fig. 5). Despite lacking these structural markers, they remain decorated by core replicase components ranging from nsp5 through the nsp8 RdRp subunit and the nsp16 methyltransferase. Their nucleic-acid-rich, round morphology is reminiscent of viral factories or ‘viroplasms’ described for other viruses^67,68,69^, but not previously reported for coronaviruses. These structures may represent separate, long-lived storage or sequestration compartments, analogous to stress granules in host cells. While RNA-protein biocondensates (speckles) are common in the nucleus^70^, it is striking that potentially related SARS-CoV-2 structures localize to the cytoplasm.

We also observe thin, linear dsRNA ‘connectors’ that bridge neighboring DMVs and sometimes tether DMV clusters to these non-canonical dsRNA granules (Fig. 4C–G, Supplementary Fig. 4A, Supplementary Fig. 5A–C). To our knowledge, these inter-organellar bridges have not been previously described for coronavirus infection. The visualization of these faint conduits was likely enabled by the high spatial resolution and optimized labeling density of our dSTORM approach, as such thin structures (∼15–20 nm) can easily be obscured by background or thresholded out compared to the extremely bright dsRNA cores in lower-resolution studies. The exact membrane topology of these connectors and granules is currently unknown. Since the massive dsRNA granules lack the canonical nsp3/nsp4 pore complex, their exact biogenesis is unclear; however, the absence of nsp3/nsp4 does not preclude the presence of a protective lipid bilayer. Considering that viral dsRNA is typically shielded from host sensors in membrane-protected environments, one compelling hypothesis is that these cytoplasmic connectors are not ‘naked’, but rather enveloped within the lumen of the host-derived ER tubules known to interconnect DMVs^42^.

Functionally, while these interconnected networks might serve as conduits for RNA traffic to seed new replication sites, their directionality cannot be inferred from our fixed-cell data. Alternatively, rather than active replication centers, these non-canonical dsRNA accumulations may represent specialized compartments for the shuttling, storage, or sequestration of excess viral dsRNA away from the primary replication sites. Recognizing that standard FISH denaturation steps are often insufficient to melt highly stable, protected dsRNA duplexes for probe annealing, definitively proving the origin of the RNA within these dense granules via hybridization is technically challenging. However, a viral origin is highly probable because: (i) comparable massive dsRNA accumulations are absent in non-infected cells, making an infection-induced burst of host dsRNA unlikely; (ii) they are specifically decorated with multiple viral replicase proteins; and (iii) they are occasionally physically linked to canonical, replication-competent DMVs via the dsRNA connectors (Fig. 4F). Defining the precise membrane topology and molecular composition of the connectors, alongside the material properties of these viroplasm-like granules, remains a key direction for future work.

While this study was not designed to fully elucidate all aspects of virion assembly and egress, the SR localization patterns of the structural proteins (Fig. 5, Supplementary Figs. 6, 7) provide critical insight. In particular, M proteins exhibit a spatial maturation trajectory, transitioning from large, unstructured conglomerates in the perinuclear region to more organized round structures partway toward the cell periphery, and finally to hollow clusters of virion-sized particles closer to the plasma membrane. The unstructured perinuclear population clearly corresponds to the Golgi bodies (Fig. 5C), and the hollow clusters correlate with lysosomal membranes in some cases. However, the M protein also displays a structured pattern in different, often large vesicles that do not contain the lysosome marker LAMP1 (Fig. 5D, blue box). SR imaging of the spike protein suggests that these structures likely represent active sites of massive virion production, storage or transport. Previous studies suggested that virions form in the ER-Golgi intermediate compartment before entering the Golgi apparatus^43,71^; however, our results suggest that M and S are processed and glycosylated in the Golgi and assemble into virions only later, possibly in both LAMP1-positive lysosomes and unidentified LAMP1-negative vesicles. This aligns with EM studies proposing virion formation in single-membrane vesicles and virion egress via “large virus-containing vesicles” that bypass the Golgi complex^72^. However, given the static nature of our imaging, we cannot definitively exclude the possibility that some of these particles correspond to mature virions re-internalized via endocytosis.

We also observe diffuse cytosolic vgRNA and N proteins that associate with each other but do not colocalize with S proteins (Fig. 5G–J), consistent with RNP formation in the cytosol independent of budding sites. Given that the N protein interacts with M in an RNA-dependent manner^73^, these RNPs likely dock onto M at budding membranes, promoting membrane closure and virion completion.

Furthermore, the lack of colocalization between S1 and S2 subunits on ER tubules and the nuclear envelope suggests that a fraction of the spike protein is cleaved and dissociates within the host cell, potentially through furin-mediated processing^74^. The extent of this S1/S2 processing represents a critical evolutionary trade-off for the virus. On one hand, cleavage at the S1/S2 boundary is advantageous and necessary for propagation, as it biases the spike protein toward an open conformation that promotes ACE2 receptor binding^75^ and enhances membrane fusion and virulence^76^. On the other hand, this cleavage intrinsically destabilizes the S protein^75^, leading to the premature intracellular shedding of S1 that we observe.

Since S1 is strictly required for viral entry, its shedding inactivates the spike protein and reduces overall infectivity, a functional cost that has been directly demonstrated in pseudotyping studies^74^. Consequently, viral evolution continuously fine-tunes this process to maximize the open conformation while limiting detrimental S1 shedding—a structural balance effectively achieved by the globally dominant D614G mutation, which stabilizes the spike to prevent shedding while efficiently binding ACE2^77^. Because our study utilized the original USA-WA1/2020 isolate, which lacks this stabilizing mutation, our spatial atlas provides a direct visualization of this pronounced, pre-adaptation shedding phenotype.

Although nirmatrelvir is clinically approved, its nanoscale effects in infected human cells have not been defined. Under M^pro^ inhibition, we frequently observe multilayered bodies (MLBs) (Fig. 6, Supplementary Fig. 9), prompting a refinement of the ER-to-DMV transformation model. The double-layered localization pattern of the nsp3 N-terminus suggests that MLBs contain zippered membranes, possibly due to the trans-interaction of nsp3 and nsp4 ectodomains^4^. Although nsp3/nsp4 expression is sufficient to generate DMVs, previous reports indicate that replacing nsp4 with an inefficiently cleaved nsp4–nsp5–nsp6 polyprotein causes ER membranes to zipper without forming DMVs^78^. We therefore propose that an uncleaved nsp4–nsp5–nsp6 polyprotein, anchored at both ends in the ER, lacks the flexibility required for cis interactions and membrane bending; cleavage between nsp4 and nsp6 is likely necessary to complete DMV formation. Given that DMV pores contain 12 copies of nsp4 in two conformations^4^, one conformation may be cleaved while the other remains uncleaved, although this requires experimental confirmation.

Together, these observations support a model in which M^pro^ cleavage acts as a molecular switch that toggles nsp3/nsp4 from trans-zippering to cis-bending, enabling DMV biogenesis (Fig. 7B). Beyond reducing RO abundance and inducing MLBs, nirmatrelvir treatment leaves residual, replication-competent DMVs, and after drug washout, infection rapidly rebounds. This suggests that partial inhibition is insufficient: effective therapy may require eliminating residual ROs and, potentially, destabilizing the polyprotein-rich MLBs defined here, to improve the durability of protease inhibition.

Although this study advances our understanding of SARS-CoV-2 infection in mammalian cells, important questions remain. A key set concerns single-stranded vgRNA once it exits replication organelles: which proteins or RNP chaperones protect and traffic it, and do RNA modifications promote its exchange to assembly sites^79^? Equally critical is defining the precise membrane topology of the novel inter-DMV connectors and nsp3/nsp4-lacking dsRNA granules identified here. Future studies must determine whether these massive accumulations represent active hubs for seeding new DMVs or specialized compartments for viral RNA storage and sequestration.

Methodologically, as shown previously^80^, correlative fluorescence–cryo-EM (CLEM) will be essential to add molecular specificity to high-resolution volumes. This approach is required to structurally confirm whether the dsRNA connectors are enveloped by the membranes of host ER tubules, and to validate the zippered membrane architecture of the nirmatrelvir-induced MLBs.

Future work should move beyond static snapshots to live-cell, single-molecule tracking. Resolving these dynamic trajectories will clarify the exact spatiotemporal sequence of polyprotein cleavage during DMV biogenesis, establish the directionality of RNA transport, and distinguish *de novo* virion assembly in LAMP1-positive vesicles from endocytic re-internalization.

The spatial organization defined here can also guide interaction studies relevant to drug design^32^. For example, given the prominence of the M protein pathway (Fig. 5A–F), disrupting M–N/vgRNA interactions is a concrete therapeutic hypothesis. More broadly, our quantitative SR imaging pipeline could be adapted for high-resolution phenotypic screening to identify novel antivirals that arrest specific spatial stages of the viral life cycle. We anticipate that the methods and maps presented here will serve as a foundational platform for dissecting vgRNA protection, trafficking, and virion creation in SARS-CoV-2 infection.

## Methods

### Antibodies

Antibodies and their working concentrations are listed in Supplementary Table 1. During the initial antibody screening, we colabeled infected cells with an anti-dsRNA antibody to evaluate the fluorescence signal in both infected (dsRNA-positive) and uninfected (dsRNA-negative) cells using DL microscopy. Antibodies were rejected if the specific signal in infected cells was too low or if the non-specific background in uninfected cells was too high. For example, validated anti-nsp1 antibodies yielded an average specific signal in late-stage infection ∼20-fold higher than the non-specific cytoplasmic signal in uninfected cells. To confirm that the fluorophore attached to the secondary antibody did not produce artifacts, we swapped the secondary antibody labels for several targets and found no difference in the resulting SR structures observed.

### Culture of cell lines

A549-ACE2 cells (human lung carcinoma cells expressing human angiotensin-converting enzyme 2; BEI Resources, NR-53821), Vero E6 cells (African green monkey kidney epithelial cells; ATCC, CRL-1586), Vero E6-Sec61β-GFP stable cells (described previously^5^), and Vero E6-TMPRSS2 cells (BPS Bioscience, 78081) were cultured in Dulbecco’s modified Eagle medium (DMEM) supplemented with GlutaMAX, 25 mM D-glucose, 1 mM sodium pyruvate (Gibco, 10569010), and 10% FBS (Sigma-Aldrich, F0926) at 37 °C and 5% CO_2_ in a humidified incubator. Cell lines were not authenticated after purchase prior to use. For A549-ACE2 cells, blasticidin (Thermo Fisher, A1113903) was added at a final concentration of 100 μg/ml. For Vero E6-TMPRSS2, geneticin (G418) was added to the culture medium at a final concentration of 1 mg/ml. Vero E6-Sec61β-GFP cells were used for Supplementary Fig. 6D; Vero E6 cells were used for Fig. 5G–J, Supplementary Figs. 5B and 8C, D. All other labeling and imaging experiments were performed using A549-ACE2 cells.

### SARS-CoV-2 viral stocks preparation

SARS-CoV-2 isolate USA-WA1/2020 (NR-52281, BEI Resources) was passaged three times in Vero E6-TMPRSS2 cells as previously described^81,82^. Briefly, a Vero E6-TMPRSS2 monolayer was infected with the virus obtained from BEI; at 72 hours post infection (hpi), P1 virus-containing tissue culture supernatants were collected and stored at −80°C. Following titration, the P1 virus stock was used to generate a P2 stock by infecting Vero E6-TMPRSS2 monolayers at a multiplicity of infection (MOI) of 0.0001 for 72 h. The P2 virus was passaged again in Vero E6-TMPRSS2 cells to obtain the P3 stock. Viral titers were determined by standard plaque assay on Vero E6 cells. All the experiments involving virus infection were performed using the P3 SARS-CoV-2 USA-WA1/2020 stock, containing a 100% wild-type (WT) population with no deletion in the spike multi-basic cleavage site.

### Infection of cells by SARS-CoV-2 (BSL-3)

Initial infections were performed under Biosafety Level 3 (BSL-3) conditions. Following the early 2025 CDC regulatory update, the safety level for SARS-CoV-2 was reduced, and subsequent experiments were performed at BSL-2; both conditions are described below as appropriate.

One day prior to infection, A549-ACE2 cells were seeded at 10,000 cells per well into 18-well glass-bottom µ-slides (Ibidi, 81817) and cultured in DMEM supplemented with 10% FBS and 100 µg/mL blasticidin. Cells were taken to the BSL-3 facility, washed once with PBS, and incubated with SARS-CoV-2 (USA-WA1/2020) at an MOI of 2 in 50 µL of DMEM containing 2% FBS for 2 h. The viral inoculum was then removed, and cells were cultured in 100 µL of DMEM containing 2% FBS for 24 h. Cells were subsequently washed with PBS, fixed with 4% paraformaldehyde (PFA; Electron Microscopy Sciences, 15710) and 0.1% glutaraldehyde (Electron Microscopy Sciences, 16350) in PBS for 1 h, and removed from the BSL-3 facility for further processing. This part of work was conducted at the high-containment BSL-3 facility of Stanford University in accordance with CDC and institutional guidelines.

### SARS-CoV-2 infection and nirmatrelvir treatment (BSL-2)

A549-ACE2 cells were seeded one day prior to infection as described above. Cells were washed once with PBS and incubated with SARS-CoV-2 WA 1 (USA-WA1/2020) at an MOI of 10 in 50 µL of DMEM containing 2% FBS for 6 h. The viral inoculum was then removed, and cells were cultured in 100 µL of DMEM containing 2% FBS and 288 nM nirmatrelvir (MedChemExpress, HY-138687) for 18 h. A subset of cells was then washed with PBS and fixed by 4% PFA and 0.1% glutaraldehyde in PBS for 1 h. For drug washout experiments, the remaining cells were washed and cultured in fresh DMEM containing 2% FBS for an additional 24 h before being washed and fixed as described above. This part of the work was conducted in a BSL-2 tissue culture room following Stanford University’s Administrative Panel on Biosafety (APB) protocols and CDC guidelines. The EC_50_ for this experiment was determined in the BSL-3 facility following a previously published protocol^83^.

### Synthesis of the RNA FISH probes

Viral genomic RNA (vgRNA) FISH probes targeting the ORF1a region of SARS-CoV-2^24^ were prepared using the same procedures as previously published^5^. RNA FISH probes targeting human 18S rRNA were designed using the Stellaris RNA FISH Probe Designer (Biosearch Technologies, version 4.2; https://www.biosearchtech.com/stellaris-designer). The 38 probe sequences are as follows: gagcgaccaaaggaaccata; accacagttatccaagtagg; tcggcatgtattagctctag; gggttggttttgatctgata; tatctagagtcaccaaagcc; gatagggcagacgttcgaat; tatttttcgtcactacctcc; cctcgaaagagtcctgtatt; tccaatggatcctcgttaaa; tcaaagtaaacgcttcgggc; attattcctagctgcggtat; acaaaatagaaccgcggtcc; ccgactttcgttcttgatta; ggtatctgatcgtcttcgaa; catcgtttatggtcggaact; tcccgtgttgagtcaaatta; cacccacggaatcgagaaag; aactaagaacggccatgcac; taaccagacaaatcgctcca; cagagtctcgttcgttatcg; gtcgcgtaactagttagcat; tgttattgctcaatctcggg; cacttactgggaattcctcg; atcaacgcaagcttatgacc; ctggcaggatcaaccaggta; tctttgagacaagcatatgc; tccggaatcgaaccctgatt; ttggatgtggtagccgtttc; ggtcgggagtgggtaatttg; gctttttaactgcagcaact; ctagcggcgcaatacgaatg; cgccggtccaagaatttcac; ggcgggtcatgggaataacg; tagggtaggcacacgctgag; caatcggtagtagcgacggg; atccgagggcctcactaaac; gatagtcaagttcgaccgtc; gttacgacttttacttcctc.

These probes were synthesized with 5’ amine-modified C6 linkers (5AmMC6) by Integrated DNA Technologies at a 25 nmol scale with standard desalting. Each probe was dissolved in nuclease-free water to a final concentration of 100 µM. Equal volumes of each probe were pooled to create a 100 µM master mix, which was subsequently desalted via ethanol precipitation. Briefly, 120 µL of the probe mix was combined with 12 µL of 3 M sodium acetate (pH 5.5) and 400 µL of ethanol. After precipitation at −80 °C overnight, probes were pelleted by centrifugation at 12,000g for 10 min at 4 °C, washed three times with precooled 70% (v/v) ethanol, air-dried, and resuspended in water to a final concentration of 100 µM. For fluorescent conjugation, 18 µL of the 100 µM probe mix was combined with 2 µL of 1 M NaHCO_3_ (pH 8.5) and 100 µg of Alexa Fluor^TM^ 647 succinimidyl ester (NHS) (Invitrogen, A37573) dissolved in 2 µL of dry DMSO (Invitrogen, D12345). The reaction was incubated for 3 days at 37 °C in the dark and purified via three rounds of cleanup using the Monarch PCR & DNA Cleanup Kit (5 µg) (NEB, T1030S) according to the manufacturer’s instructions.

### RNA FISH and immunofluorescence (IF)

RNA FISH and IF staining were performed as previously described^5^.

### Western blotting

A549-ACE2 cells were seeded at a density of 2.8 × 10⁵ cells per well in 6-well plates and cultured in DMEM supplemented with 10% FBS and 100 µg/mL blasticidin. The following day, cells were infected with SARS-CoV-2 at a MOI of 10. After a 6-hour incubation at 37 °C, the viral inoculum was removed, and cells were treated for 18 hours with either 288 nM nirmatrelvir or an equivalent volume of DMSO control in DMEM with 2% FBS. A separate well of uninfected cells was maintained as a negative control. At 24 hours post-infection, cells were washed three times with PBS and lysed on ice for 30 minutes in RIPA Lysis and Extraction Buffer (Thermo, 89900) supplemented with 1x cOmplete™ Protease Inhibitor Cocktail (Sigma, 11836170001). Cell debris was pelleted by centrifuging lysates at 16,000 × g for 20 minutes at 4°C. The protein concentration of the supernatant was determined using a Pierce™ BCA Protein Assay Kit (Thermo, A65453).

6 µg of protein samples were reduced and denatured in 4x NuPAGE™ LDS Sample Buffer (Thermo, NP0007) with DTT by heating at 70°C for 10 minutes, then resolved by gel electrophoresis (Thermo, NP0321BOX). The HiMark™ Pre-stained Protein Standard (Thermo, LC5699) and PageRuler™ Plus Prestained Protein Ladder (Thermo, 26619) were included as molecular weight markers. Proteins were transferred to PVDF membranes. For nsp3 detection, a wet tank transfer was used, while for ACTB, the iBlot 2 Dry Blotting System (Thermo Fisher Scientific) was employed. Membranes were blocked for 1 hour at room temperature in 5% BSA in Tris-buffered saline (Thermo, J60764.K2) containing 0.1% Tween-20 (TBST).

For nsp3, membrane was incubated overnight with gentle rocking at 4°C with the rabbit anti-nsp3 primary antibodies (1:2,000; Thermo, PA5-116947), diluted in blocking buffer. After washing three times with TBST, membrane was incubated with HRP-conjugated goat anti-rabbit IgG (H+L) secondary antibody (1:5,000; Thermo, A16035) for 30 minutes at room temperature. For ACTB, membrane was incubated with Direct-Blot™ HRP-conjugated anti-β-actin antibody (1:1,000; BioLegend, 643808) with gentle rocking for 30 minutes at room temperature. Membranes were washed with 1x TBST buffer three times. Following final washes, bands were visualized using SuperSignal™ West Pico PLUS Chemiluminescent Substrate (Thermo, 34577) and imaged using an Azure Imaging System (Azure Biosystems 300). For a subset of biological replicates, an alternative fluorescent detection method was used for nsp3. Following incubation with the primary antibody as described above, membranes were incubated with IRDye® 800CW donkey anti-rabbit IgG secondary antibody (1:5,000; Licorbio, 926-32213) for 30 minutes at room temperature. The membrane was washed three times with 1x TBST buffer and imaged using a LI-COR Odyssey imaging system and the Image Studio 6.0 software.

### Spinning-disk confocal microscopy

Confocal microscopy was performed at the Stanford University Cell Sciences Imaging Core Facility using a Nikon Ti2 Crest spinning disk confocal (SDC) microscope equipped with a Photometrics Kinetix camera, a Perfect Focus System (PFS), and four laser lines (365, 488, 561, and 640 nm). Images were acquired through a 50 µm pinhole using either a 20× Plan Apo air objective (NA = 0.80; 0.333 µm/pixel) or a 60× Plan Apo oil objective (NA = 1.40; 0.108 µm/pixel). Z-stacks were captured at a 0.3 µm step size using a piezo Z-axis stage (Mad City Labs) and the NIS Elements software (v5.42.06).

### Confocal image analysis

Image processing was performed in Fiji (ImageJ 1.54p). Projections of adjacent Z planes exhibiting maximum loci fluorescence were generated. For conditions 1 and 2 (Supplementary Fig. 9A), both infected and total cells were counted manually using the Fiji multi-point tool. For condition 3, which contained several hundred cells per field of view, infected cells were counted manually, while total cells were quantified using an automated segmentation pipeline. Briefly, nsp4 and nsp5 fluorescence images were smoothed with a Gaussian blur (*σ* = 3.5 pixels), thresholded, and binarized. Intracellular holes were filled, and total cell counts were obtained via particle analysis with a size threshold of >30 µm^2^. The infection ratio was calculated as the number of infected cells divided by the total number of cells in each condition. Statistical analysis was performed via one-way ANOVA using GraphPad Prism.

### Classification of Infection Stages

While all samples were fixed at a single chronological timepoint (24 hpi), the infection progression in A549-ACE2 cells is asynchronous. We classified cells into ‘Early’ or ‘Late’ infection stages based on the distinct fluorescence morphology of the replication organelles^5^. ‘Early stage’ cells were defined by the presence of distinct, individual DMVs (typically ≲ 200 nm in diameter) distributed in the cytoplasm, while ‘Late stage’ cells were identified by the presence of large, merged vesicle packets and dense perinuclear accumulations. This morphological transition is consistent across various viral targets (Supplementary Fig. 1). For S and M labeling, ‘Early stage’ was defined by the presence of fluorescence signal in the cytoplasm with a minimal amount of virion-like particles at the cell surface, and the ‘Late stage’ was defined by the presence of a high density of virus particles at the cell surface. Cells with ambiguous morphology were excluded from stage-specific analysis.

### Optical setup for two- and three-color 2D SR microscopy

Two- and three-color 2D (d)STORM SR imaging was performed on a custom-built microscope^5^ based on a Nikon Diaphot 200 inverted frame equipped with a 60x/1.35 NA oil-immersion objective (Olympus UPLSAPO60XO) and an electron-multiplying Si CCD (EMCCD) camera (Andor iXon Ultra 897, 512 × 512 pixels). Samples were mounted on stacked coarse (U-780.DOS) and fine (P-545.3C8S) piezo stages (Physik Instrumente). Fluorophore excitation was driven by 642 nm and 560 nm continuous-wave (CW) lasers (1 W each; MPB Communications) for AF647/CF680 and CF583R/CF568, respectively. A 405 nm CW diode laser (50 mW; Coherent OBIS) was used for fluorophore reactivation. Excitation power was modulated by adjusting the laser output and, when necessary, via neutral density filters housed in a motorized filter wheel (FW102C, Thorlabs). All laser beams were coupled into a square-core multimode fiber equipped with a shaker for speckle reduction (Newport F-DS-ASQR200-FC/PC). The fiber output face (200 × 200 µm^2^ core) was imaged with a 10×/0.25 NA objective and magnified to achieve uniform, flat-field illumination over a 47.6 × 47.6 µm^2^ region in the sample plane.

Fluorescence emission was separated from the excitation light using a multiband dichroic mirror (ZT405/488/561/640rpcv2, Chroma) and subsequently divided into two spectral channels at λ ≈ 690 nm with a dichroic mirror (T690LPxxr, Chroma). The long-wavelength channel (λ > 690 nm) was filtered with an ET700/75M bandpass (Chroma) and used for simultaneous ratiometric imaging of CF680 and AF647. The short-wavelength channel (λ < 690 nm) served a dual purpose: it functioned as the second channel for CF680/AF647 ratiometry (filtered with a ZET635NF notch and ET685/70M bandpass, Chroma), or as a dedicated channel for CF583R imaging (filtered with a 561LP long-pass and 607/70BP bandpass, Semrock). For rapid remote filter transition, these short-wavelength filters were housed in a second motorized filter wheel (FW102C, Thorlabs). Images were formed with a 400 mm tube lens and relayed via 120 mm lenses, projecting both channels side-by-side onto the camera chip with an effective pixel size of 117 × 117 nm^2^ in the sample coordinates.

Axial drift was actively compensated using a custom focus lock system based on an infrared beam reflected from the coverslip-sample interface, as previously described^5^. The microscope was operated via custom MATLAB R2023b scripts (MathWorks) that integrated control of the focus lock, lasers, shutters, filter wheels, and the fine piezo stage. EMCCD camera data acquisition was controlled via Andor SOLIS software (version 4.31.30024.0).

### SR imaging procedure

For (d)STORM, each well of the 18-well chamber was filled with 200 µl of photoblinking buffer, consisting of 200 U/ml glucose oxidase, 1000 U/ml catalase, 10% (w/v) glucose, 200 mM Tris-HCl (pH 8.0), 15 mM NaCl, and 50 mM cysteamine. The buffer was prepared fresh by mixing three stock solutions^20^: (1) an enzyme mix containing 4000 U/ml glucose oxidase (G2133, Sigma), 20,000 U/ml catalase (C1345, Sigma), 25 mM KCl (P217, Fisher), 4 mM TCEP (646547, Sigma), 50% v/v glycerol (BP229, Fisher), and 22 mM Tris-HCl (pH 7.0) (BP1756, Fisher), stored at −20 °C; (2) 1 M cysteamine-HCl (30080, Sigma), stored at −20 °C; and (3) a substrate stock containing 37% w/v glucose (49139, Sigma), 56 mM NaCl (S271, Fisher), and 0.74 M Tris-HCl (pH 8.0) (J22638.AE, Fisher), stored at 4 °C.

For samples incorporating RNA FISH, the buffer was supplemented with 1 U/µl of an RNase inhibitor (302811, LGC Biosearch Technologies) and the bovine liver catalase was replaced with catalase from *Aspergillus niger* (219261, Sigma), as this was found to be less contaminated with RNases. To prevent photobleaching in non-imaged adjacent wells, they were maintained in an oxygen-scavenging buffer containing 100 U/ml glucose oxidase, 500 U/ml catalase, 4.6% (w/v) glucose, 93 mM Tris-HCl (pH 8.0), and 7 mM NaCl. After removing the imaging or oxygen scavenging buffer, samples were stored in PBS supplemented with 0.02% sodium azide (cs-296028, ChemCruz) at 4 °C to prevent bacterial growth.

Before SR imaging, the region of interest (ROI) was selected using standard wide-field diffraction-limited (DL) fluorescence. For three-color (d)STORM experiments, we first simultaneously imaged CF680 and AF647 in the two spectral channels using 642 nm excitation (power density ∼20 kW/cm^2^), and then switched to CF583R at ∼13 kW/cm^2^ of 560 nm excitation. For two-color (d)STORM, we first imaged AF647 using only the short-wavelength (λ < 690 nm) channel at 9–20 kW/cm^2^ of 642 nm, and then CF583R at ∼13 kW/cm^2^ of 560 nm. When the single-molecule (SM) blinking density was low, the sample was additionally illuminated with 405 nm light (up to 50 W/cm^2^) for faster fluorophore reactivation. We used an exposure time of 10.57 ms per frame and an EM gain of 84. Image acquisition commenced after the initial dark state shelving phase, once clear SM blinking was established; movies were acquired for approximately 8 × 10^4^ frames for each fluorophore imaged separately, or for ∼1.6 × 10^5^ frames when CF680 and AF647 were imaged simultaneously.

### SR data analysis

SM movies were first processed using the ThunderStorm (dev-2016-09-10-b1) plugin^84,85^ for Fiji^86^ with the following parameters: image filtering via wavelet filter (B-spline order = 3, scale = 2); approximate localization using a local maximum search (threshold = *A* × std(Wave.F1), 8-neighbourhood connectivity); and sub-pixel localization using an integrated Gaussian fitting model (fitting radius = 4 pixels, weighted least squares method, initial *σ* = 1.1 pixels). The threshold multiplier *A* was set to 0.9 for simultaneously imaged CF680 with AF647 in both spectral channels, and to 1.5−2.0 for fluorophores imaged separately.

After fitting, data for separately imaged fluorophores were corrected for drift using cross-correlation in ThunderStorm and filtered to retain localizations with a fitted Gaussian width (*σ*) <200 nm. If ThunderStorm drift correction failed, these datasets were corrected using SharpViSu (v2.0.3)^87^. For simultaneously imaged CF680 and AF647, drift was not corrected at this stage; localizations from both channels were first filtered (80 nm < *σ* < 200 nm), and duplicates within a 100 nm radius were removed. These dual-channel localizations were then demixed in SplitViSu^20^. Because drift is identical across simultaneously imaged channels, it was corrected in SharpViSu using cross-correlation between temporal subsets of the combined CF680 and AF647 localizations ^87^.

For further processing, we kept only localizations with fitted Gaussian *σ* between 80 nm and 160 nm to reject out-of-focus signals. Localizations found within 50 nm on consecutive frames, likely originating from multiple re-localizations of the same molecule, were processed in two ways. For SR image reconstruction, to improve resolution, these localizations were refined by selecting them from a normal distribution with a mean equal to the weighted mean of the initial localizations (weights *w_i_* =√(*N_ph,i_*)) and a standard deviation (SD) equal *σ_0_*(∑*N_ph,i_*) ^−1/2^, where *σ_0_* is the SD of the localization position in the given consecutive series and *N*_ph,i_ is the number of photons acquired from the *i*-th localization in the series^20^. For quantitative spatial analysis (to reduce overcounting), the localizations of the consecutive series were reduced to a single localization at the weighted mean position. After this correction, to suppress spurious sparse background localizations originating from photoblued far-red fluorophores^88^ or from non-specific binding, the SR data for CF583R were additionally filtered by removing localizations that had ≤4 neighbors within a 30 nm radius.

For image registration, we imaged 200 nm TetraSpeck beads (T7280, Thermo Fisher Scientific) in both the far-red and the CF583R channels and localized the beads by fitting using ThunderStorm as described above. The transformation between channels was calculated using an affine transformation matrix via the MATLAB function ‘fitgeotrans’. The calculated transformation was then applied to the CF583R localizations using the MATLAB function ‘transformPointsInverse’. After all corrections, final SR images were reconstructed as 2D histograms with a pixel size of 20 × 20 nm^2^ (typically used for whole-cell images) or 16 × 16 nm^2^ (typically used for zoomed-in images).

### 3D DHPSF imaging

Two-color 3D microscopy was performed using a previously described, custom-built 3D SM imaging microscope^89^. Briefly, 647 nm, 561 nm, and 405 nm lasers were co-aligned and focused onto the fiber input of a multimode fiber de-speckler (Newport). The fiber output was imaged into the sample plane by reflecting off a quad-band dichroic mirror and passing through a 100×, 1.4NA oil-immersion objective lens, producing a 35-μm diameter circular illumination pattern at the sample. Collected fluorescence emission passed through the quad-band dichroic and was relayed to the camera via a two-channel *4f* system, with a 660 nm long-pass dichroic mirror splitting the emission. Each arm of the emission pathway was equipped with a double-helix phase mask^90^ (Double-Helix Optics) placed in the Fourier plane to produce the double-helix point-spread function (DHPSF) response in each channel on the detector. A knife-edge prism mirror was used to direct each channel onto opposite quadrants of an EMCCD. Samples were first imaged with 647 nm illumination (7.5 kW/cm^2^; 17 ms exposure time; EM gain 272), followed by 561 nm illumination (6.9 kW/cm^2^; 15 ms exposure; EM gain 272). In both cases, 405 nm illumination was gradually increased from 0.6 W/cm^2^ to 60 W/cm^2^ to photoactivate molecules as emitter density decreased.

After cellular imaging, 100 nm TetraSpeck beads (Invitrogen, T7279) suspended in 1% (w/v) agarose in water were imaged for calibration. First, the bead sample was laterally scanned to image ∼3000 beads evenly distributed throughout the imaging volume. A Z-scan (3.2 μm range; 50 nm step size) for a single bead in the center of the field of view was then acquired simultaneously in each channel to compute a calibration mapping the DHPSF lobe angle to the Z-coordinate.

Single-molecule DHPSF images were analyzed with a plug-in^91^ for the PYthon Microscopy Environment (PYME 23.06.15)^92^. For each frame, the background was estimated using the median of the previous 30 frames and then subtracted. Candidate DHPSF ROIs for fitting were then extracted via a steerable-filter-based algorithm^91^. Each candidate ROI was then fitted to a DHPSF model function using least-squares optimization. Fitting parameters included lobe width (*σ*), lobe separation, individual lobe amplitudes, local background, and lobe angle (which was subsequently mapped to Z using the calibration data). After localizing cellular data, TetraSpeck bead data were localized and used to compute a 3D locally weighted mean quadratic transformation to map 561 nm channel localizations to the 647 nm channel. Localizations were then registered, drift-corrected via cross correlation, and scaled in Z by a factor of 0.7 to account for the focal shift caused by the refractive index mismatch between the aqueous medium and the cover glass. Localizations from the 647 nm channel were then filtered, retaining lobe widths of 140–275 nm and lobe separations of 900–1400 nm. Similarly, 561 nm channel localizations were filtered to retain lobe widths of 125–245 nm and lobe separations of 700–1200 nm. 3D localizations were rendered as pointsprites using PYMEVisualize^93^.

### Assessment of localization precision

For 2D (d)STORM imaging, the lateral localization precision (σ_xy_) was quantified from the single-molecule fitting data using ThunderSTORM. Based on the photon counts and local background of the emission events, the median theoretical localization precision was estimated to be ∼13 nm for AF647 and ∼17 nm for CF583R. For 3D-DHPSF imaging, the localization precision was assessed using the Cramér-Rao lower bound (CRLB) of the DHPSF fitting model in the PYME plugin^91^. The estimated median precisions for the 3D datasets were approximately 18 nm laterally (σ_xy_) and 29 nm axially (σ_z_) for AF647, and 26 nm laterally and 45 nm axially for CF583R.

### Bivariate pair-correlation functions

Bivariate pair-correlation functions, *g_12_*(*r*)^94^, were calculated by counting the number of second-species localizations situated at a distance between *r* and *r* + *dr* from each first-species localization^5^. These counts were then normalized by the area of the corresponding annulus and the average density of the second species within the ROI. The normalized values were then averaged across all first-species localizations. The radial distance, *r* was evaluated from 0 to 500 nm using a step size (*dr*) of 1 nm. To establish a baseline for complete spatial randomness (CSR), simulated datasets were generated matching the average experimental density within the same ROI, and *g_12_*(*r*) traces for these simulated CSR datasets were computed identically. Edge-effect corrections were not applied, leading to a slight, expected decrease of *g_12_*(*r*) at large *r*. In the accompanying figures, single-cell experimental and CSR *g_12_*(*r*) traces are displayed as faint lines, with the sample mean denoted by a bold line.

### Radial density distribution functions for three-color SR images of DMVs

Radial density distributions were evaluated for each dsRNA cluster using the centroid position of dsRNA as the origin. First, a ROI was manually drawn in an SR image to select DMV-like dsRNA clusters surrounded by nsp4 localizations. Irregularly shaped dsRNA clusters or those lacking nsp4 were excluded. The dsRNA centroid positions were calculated by fitting the dsRNA SR images (pixel size = 20 × 20 nm^2^) in ThunderStorm using the following parameters: image filtering via wavelet filter (B-spline order = 4, scale = 4); approximate localization using a local maximum search (threshold = 0.5 std(Wave.F1), 8-neighbourhood connectivity); and sub-pixel localization using an integrated Gaussian fitting model (fitting radius = 5 pixels, weighted least squares method, initial *σ* = 2 pixels).

For datasets containing vgRNA, fitting was performed on images combining both dsRNA and vgRNA. The SR images of both vgRNA and dsRNA were first smoothened with a Gaussian filter (*σ* = 0.5 pixels), and the combined image was calculated as the sum of pixel values from the two filtered SR images. After fitting, the dsRNA centroid localizations were filtered by discarding localizations closer than 3*σ*, and retaining only those with a fitted *σ* > 15 nm and containing >200 SM localizations.

For each dsRNA centroid *i* under a given labeling condition, the number of localizations of the three co-imaged targets within a distance between *r* and *r* + *dr* (where *dr* = 1 nm) from the centroid was counted. These counts were then normalized by the area of the corresponding annulus, yielding a distribution proportional to the line profile of the SR image passing through the dsRNA centroid, averaged across all angles. The radial distributions for each centroid *i* were smoothed using the MATLAB ‘smoothdata’ function (Gaussian method, window size = [6 6]). The resulting radial distribution curves were normalized such that the peak of their sum equaled unity, and the individual curves were multiplied by *n* (the total number of analyzed clusters in the dataset) to yield *g_i_*(*r*).

The final radial density distribution function, *g*(*r*), was calculated as the mean of all individual curves *g_i_*(*r*). For each radial position *r*, the 95% confidence interval (CI) of the mean was calculated as *g*(*r*) ± 1.96 SD[*g_i_*(*r*)] / √*n*, where SD[*g_i_*(*r*)] is the standard deviation across the normalized and rescaled curves of all clusters *i* at the given *r*.

To determine the peak position *R* of *g*(*r*), a Gaussian function with a baseline offset was fitted to *g*(*r*) in the vicinity of the peak This fit typically incorporated values down to ∼0.9*g*(*R*) on the left side of the peak and ∼0.6*g*(*R*) on the right side, and was visually inspected for accuracy. The precision of the peak fitting was evaluated using bootstrapping: the individual *g_i_*(*r*) traces were randomly resampled with replacement 1000 times. The peak positions of the resampled mean distributions, *g_j_*(*r*) (*j* = 1…1000), were fitted as described above. The standard error (SE) of the peak position was calculated as the SD of these 1000 bootstrap peaks (*R_j_*), and the 95% CI of the peak was expressed as *R* ± 1.96 SD(*R_j_*).

### Bivariate angular pair-correlation functions

Bivariate angular pair-correlation functions, *p*(*θ*) were calculated using the same ROIs as for *g*(*r*). For each dsRNA centroid, only localizations with a radial distance from the closest dsRNA centroid *r* bounded by 0.5*σ*_dsRNA_ < *r* < 4*σ_dsRNA_* were considered, where *σ*_dsRNA_ is the fitted width of the dsRNA cluster derived from ThunderStorm as described above. Localization angles were calculated using the two-argument arctangent function (atan2), setting the dsRNA centroid as the origin of the polar coordinate system. The angular separation between all nsp4 localizations and those of the second target were calculated for every dsRNA cluster. The function *p*(*θ*) was estimated by constructing a histogram of these angular separations with a bin width of π/100 radians, and was normalized such that *p*(*θ*) ≈ 1 for uncorrelated localizations.

### Theoretical Model of DMV Projection

To quantify the radial distribution of proteins, we modeled the double-membrane vesicles (DMVs) as spherical shells of true geometric radius *R*_geo_, uniformly decorated with antibody-labeled proteins. In the limit of infinite localization precision, the ideal 2D projected density *P*_ideal_(*r*) of a spherical shell is given by the Abel transform of the 3D surface distribution *δ*(*ρ−R*_geo_):

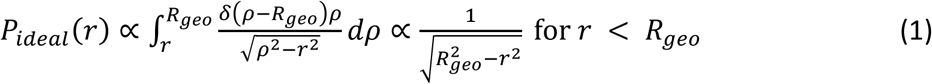

where *r* is the radial distance from the center of the projected image. This geometric projection creates a sharp intensity singularity at the edge (*r* = *R*_geo_).

To account for the finite localization precision of the SMLM system and the linkage error of the antibodies, we modeled the effective Point Spread Function (PSF) of the projected image. Since the radial profile is derived from the 2D projection of the DMVs (integration along the optical axis), the axial elongation of the PSF integrates out. Furthermore, because the final profile is obtained by averaging a large population of DMVs distributed at random axial positions, depth-of-field effects and z-dependent PSF variations are also averaged away. Consequently, the relevant blur kernel is determined solely by the effective lateral localization precision, modeled as a Gaussian with SD *σ_xy_*.

The theoretically observed radial density distribution function *g*(*r*) is the convolution of the ideal geometric projection with this lateral Gaussian PSF:

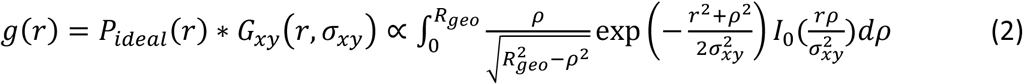

where *I_0_* is the modified Bessel function of the first kind.

To experimentally determine the radius of each protein layer, we extracted the experimental *g(r)* from averaged DMVs and fitted the peak position (*r*_peak_) using a Gaussian function with a baseline offset to account for non-specific background localizations, which provides a good approximation of the intensity maximum.

We performed numerical simulations to assess the accuracy of using the peak position *r*_peak_ as a proxy for the true radius *R*_geo_. Inspection of the integral (2) reveals that by introducing dimensionless variables scaled by the localization precision (*ȓ* = *r*/*σ*_*x,y*_, ρ̂ = ρ/*σ*_*x,y*_, *R̂* = *R_geo_*/*σ*_*x,y*_), the profile shape depends exclusively on the dimensionless radius *R̂*. Consequently, the normalized systematic peak shift (*R*_geo_ − *r*_peak_)/*σ_xy_* is a universal function of the sphere’s curvature *R*_geo_/*σ_xy_*.

To quantify this, we analyzed the projection in the “thin shell limit” (*R*_geo_ ≫ *σ_xy_*). In this regime, the geometric projection near the edge (*r* ≈ *R*_geo_) allows the term under the square root to be approximated as 2*R_geo_*(*R_geo_* − *r*), leading to 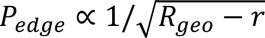. Furthermore, the integration domain can be extended to infinity, treating the edge as an isolated semi-infinite boundary. The condition for the intensity maximum 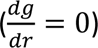 then reduces to a universal integral equation independent of *R_geo_*:

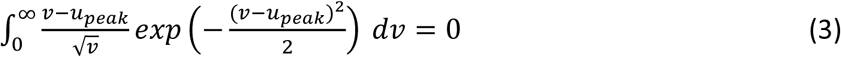

where *u* = (*R*_geo_ − *r*)/*σ*_xy_ is the dimensionless distance from the edge. Numerical solution of this equation yields a constant shift factor *u*_peak_ ≈ 0.77. Thus, in the limit of negligible curvature, the peak shift Δ = *R*_geo_ − *r*_peak_ is linear with the localization precision, and the true radius is

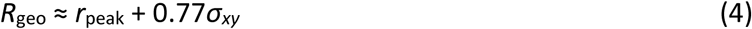

For our experimental geometry (*R*_geo_ ≈ 100 nm*, σ_xy_* ≈ 20 nm), curvature effects slightly increase this shift to approximately 0.9*σ_xy_* (Supplementary Fig. 3).

While the theoretical shift Δ is well-defined for a single particle, the averaging of DMVs with variable radii introduces an additional broadening term to the effective PSF. This population variability decreases the effective ratio *R*_geo_/*σ_xy_* and consequently increases the magnitude of the inward peak shift by an unknown amount. Since the exact radius distribution of the DMV population cannot be precisely deconvolved, correcting for the absolute geometric radius *R_geo_* would introduce model-dependent uncertainties. Therefore, throughout the figures and text, we report the directly measured peak position *r*_peak_ (denoted simply as Radius or R). However, as the systematic shift affects concentric layers with similar *R*_geo_ and *σ_xy_* nearly equally, it effectively cancels out in the relative distance measurement. Consequently, the relative radial distance Δ*R* = *R*(*i*) − *R*(*j*) provides a robust approximation of the true physical separation between the layers of proteins *i* and *j*, largely independent of the absolute peak shift.

### Statistics and Reproducibility

The experimental measurements were replicated at least 2–3 times for each labeling combination, representing independent biological replicates starting with cell culture and viral infection. Following initial protocol optimization, all subsequent biological replicates were successful. Each sample preparation yielded multiple imaging chambers containing up to 18 wells with distinct target combinations. For each sample (each target combination), typically 5–15 cells were imaged via SR microscopy. With certain targets appearing in many combinations of labels, they were independently imaged multiple times across the study. The total number of cells imaged for each primary antibody is detailed in Supplementary Table 1.

## Data availability

The Atlas described in this paper is provided as high-resolution fluorescence SR images of each infected cell at the Stanford Data Repository, which can be freely accessed at this DOI: https://doi.org/10.25740/my248kj2080. This also includes SR images of the accessory proteins ORF3a, ORF6, ORF7a, ORF8 and ORF9b which are provided as an additional resource. Source Data for the plots in the figures are provided with this paper.

## Code availability

Custom code used for image and data analysis is available in the Stanford Data Repository under the same identifier: https://doi.org/10.25740/my248kj2080

## Acknowledgements

We thank the Stanford In vitro BSL3 Service Center, its Director Jaishree Garhyan, and Amol Pohane for assistance with this research. We also acknowledge Stanford University Cell Sciences Imaging Core Facility (RRID:SCR_017787). The following reagents were obtained through BEI Resources, NIAID, NIH: Human Lung Carcinoma Cells Expressing Human Angiotensin-Converting Enzyme 2 (A549-ACE2), NR-53821, and SARS-Related Coronavirus 2, Isolate hCoV-19/USA-WA1/2020, NR-52281.

## Author Contributions Statement

L.A., M.H., L.S.Q. and W.E.M. conceived the project. L.A. designed the optical setup, performed SR imaging and data analysis. M.H. performed cell culture, viral infections, nirmatrelvir treatments, and labeling. Y.Z. performed confocal imaging and data analysis, and helped with sample preparation and with SR data analysis. A.B. performed 3D DHPSF SR imaging and analysis. L.A. and W.E.M. wrote the manuscript with input from all authors.

---

**Supplementary Fig. 1:**
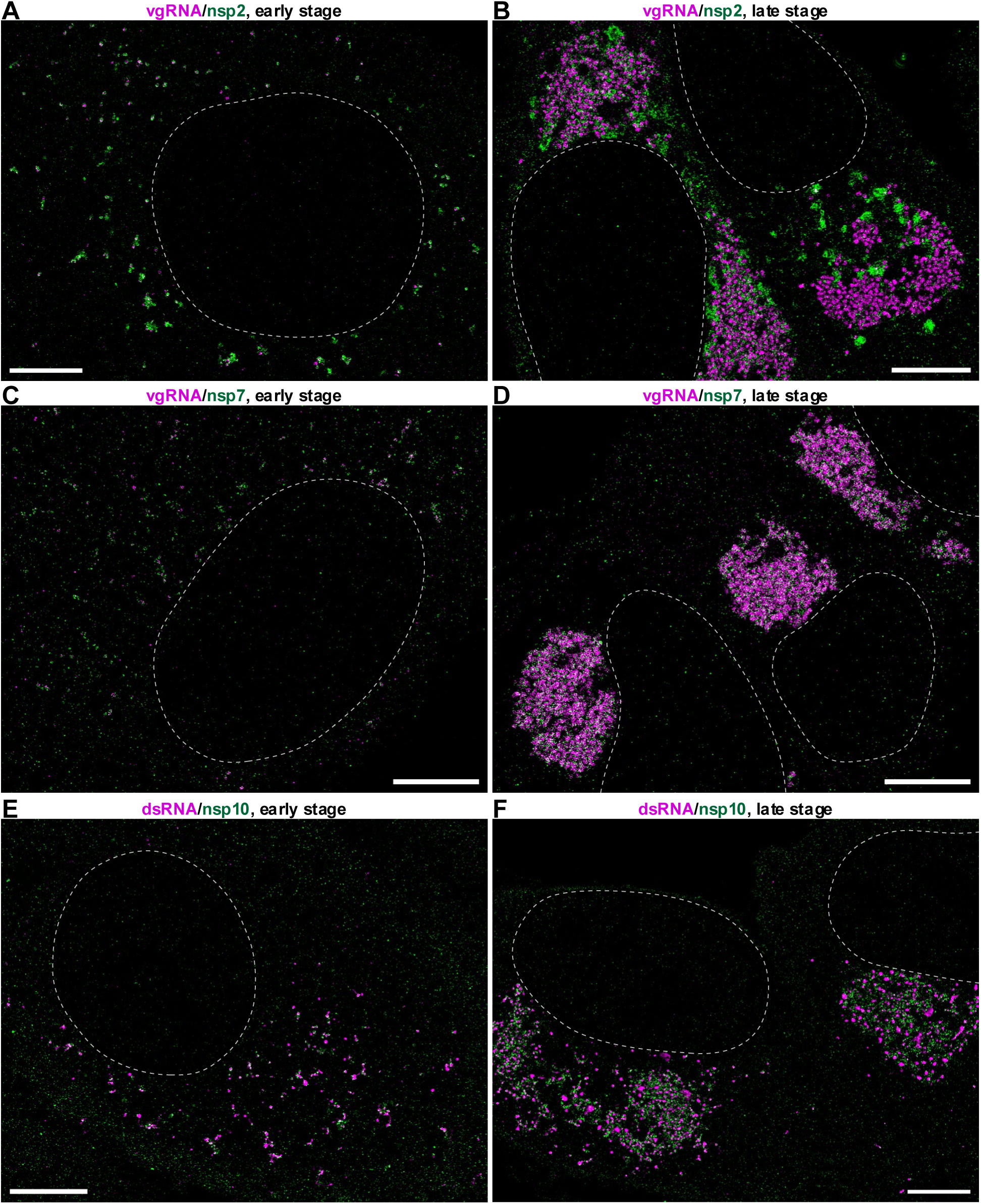
Morphological classification of infection stages in asynchronous cell populations. Although all samples were fixed at 24 hours post-infection (hpi), the viral life cycle in A549-ACE2 cells is asynchronous, resulting in a population containing cells at different functional phases of replication. Cells were stratified based on the morphology of replication organelles (ROs) observed via super-resolution microscopy. (**A**, **C**, **E**) “Early morphological stage” is characterized by isolated, individual DMVs (typically ≲ 200 nm in diameter) distributed in the cytoplasm. (**B**, **D**, **F**) “Late morphological stage” is defined by the presence of large, merged vesicle packets and dense perinuclear accumulations of viral content. This morphological transition is consistent across different replicase components and RNA species, as shown by side-by-side comparisons of vgRNA (magenta) and nsp2 (green) (**A**, **B**); vgRNA (magenta) and nsp7 (green) (**C**, **D**); and dsRNA (magenta) and nsp10 (green) (**E**, **F**). The white dashed curves indicate the approximate position of the nucleus edge. Scale bars: 5 µm.

**Supplementary Fig. 2:**
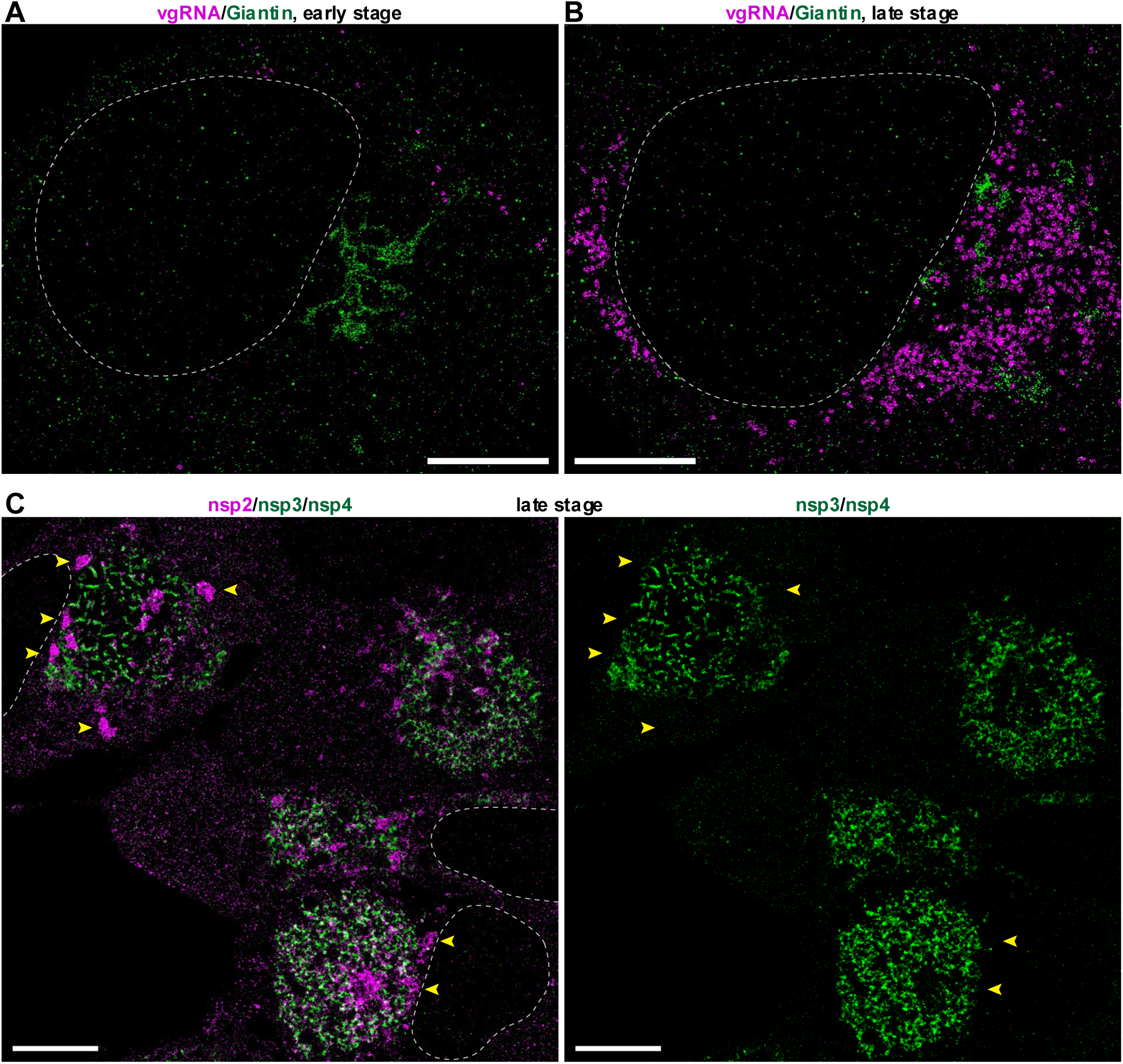
The Golgi marker Giantin remains spatially distinct from replication organelles, and nsp2 segregates from nsp3/nsp4 during Golgi remodeling. **(A)** SR image of vgRNA (magenta) and Giantin (green) in the early morphological stage indicates that the Golgi apparatus retains its typical compact, perinuclear ribbon-like structure and is spatially distinct from the isolated vgRNA puncta−early ROs. **(B)** In the late morphological stage, the Golgi apparatus undergoes fragmentation and dispersal. The Giantin signal remains separate from the large, merged vgRNA clusters, indicating that Giantin is not relocalized to or recruited into the viral replication organelles. **(C)** SR image of nsp2 (magenta) costained with a mixture of anti-nsp3 and nsp4 antibodies (green) in the late infection state. Nsp2 accumulates in dispersed structures consistent with remodeled Golgi bodies (yellow arrowheads) which are distinctly devoid of nsp3 and nsp4 signals. The white dashed curves approximate the position of the nuclear edge. Scale bars, 5 µm.

**Supplementary Fig. 3.**
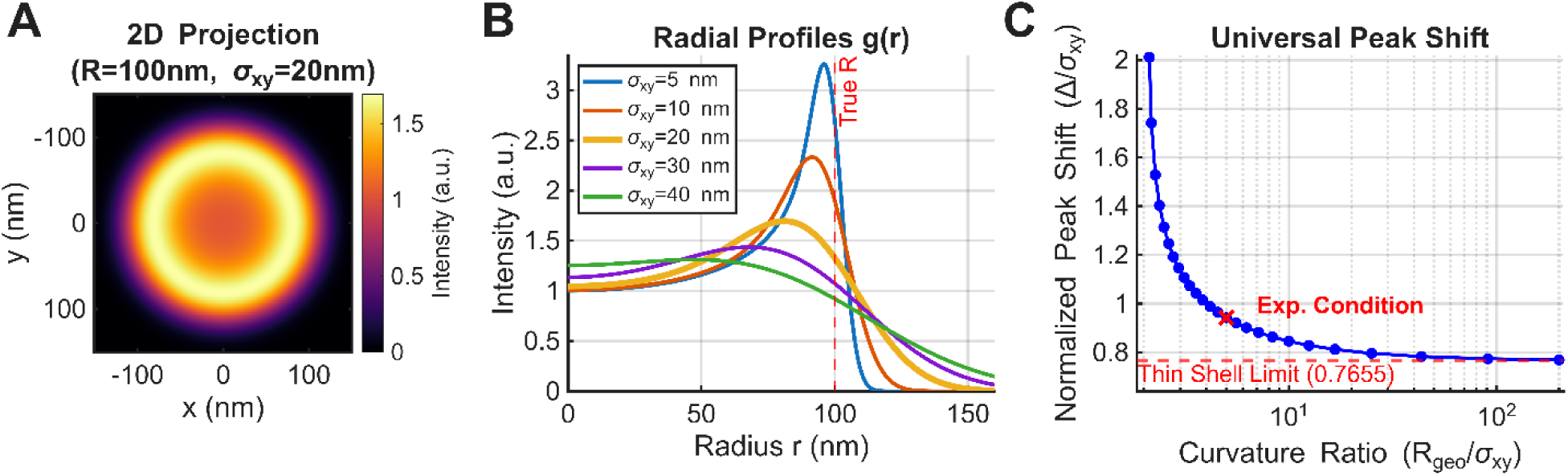
Simulation of projection effects and systematic peak shift in DMV radius measurements. **(A)** Simulated 2D projection of a double-membrane vesicle (DMV) modeled as a spherical shell with geometric radius *R*_geo_ *=* 100 nm and lateral localization precision *σ_xy_ =* 20 nm. The image represents the convolution of the ideal geometric projection (Abel transform) with a Gaussian Point Spread Function (PSF). **(B)** Theoretical radial intensity profiles *g*(*r*) calculated for a fixed geometric radius (*R_geo_ =* 100 nm) and varying localization precisions (*σ_xy_ =* 5−40 nm). The profile calculated for *σ_xy_ =* 20 nm (bold yellow curve) strongly resembles the experimental profiles for many DMV-associated proteins, with the exception of an additional nearly constant background that is trivial and was neglected in this theoretical development. The vertical dashed line indicates the true geometric radius. Note that as the blur width (*σ_xy_*) increases relative to the shell radius, the position of the intensity maximum (*r*_peak_) shifts systematically inward. **(C)** Universal master curve quantifying the systematic peak shift Δ = *R*_geo_ − *r*_peak_. The shift, normalized by the localization precision (Δ/*σ_xy_*), is plotted as a function of the dimensionless curvature ratio (*R*_geo_/*σ_xy_*). In the thin-shell limit (*R*_geo_ ≫ *σ_xy_*), the shift approaches a constant value of ≈ 0.77*σ_xy_* (dashed red line). For the experimental conditions (indicated by the red cross; *R_geo_* ≈ 100 nm, *σ_xy_* ≈ 20 nm), curvature effects slightly increase this shift to approximately 0.9*σ_xy_*.

**Supplementary Fig. 4:**
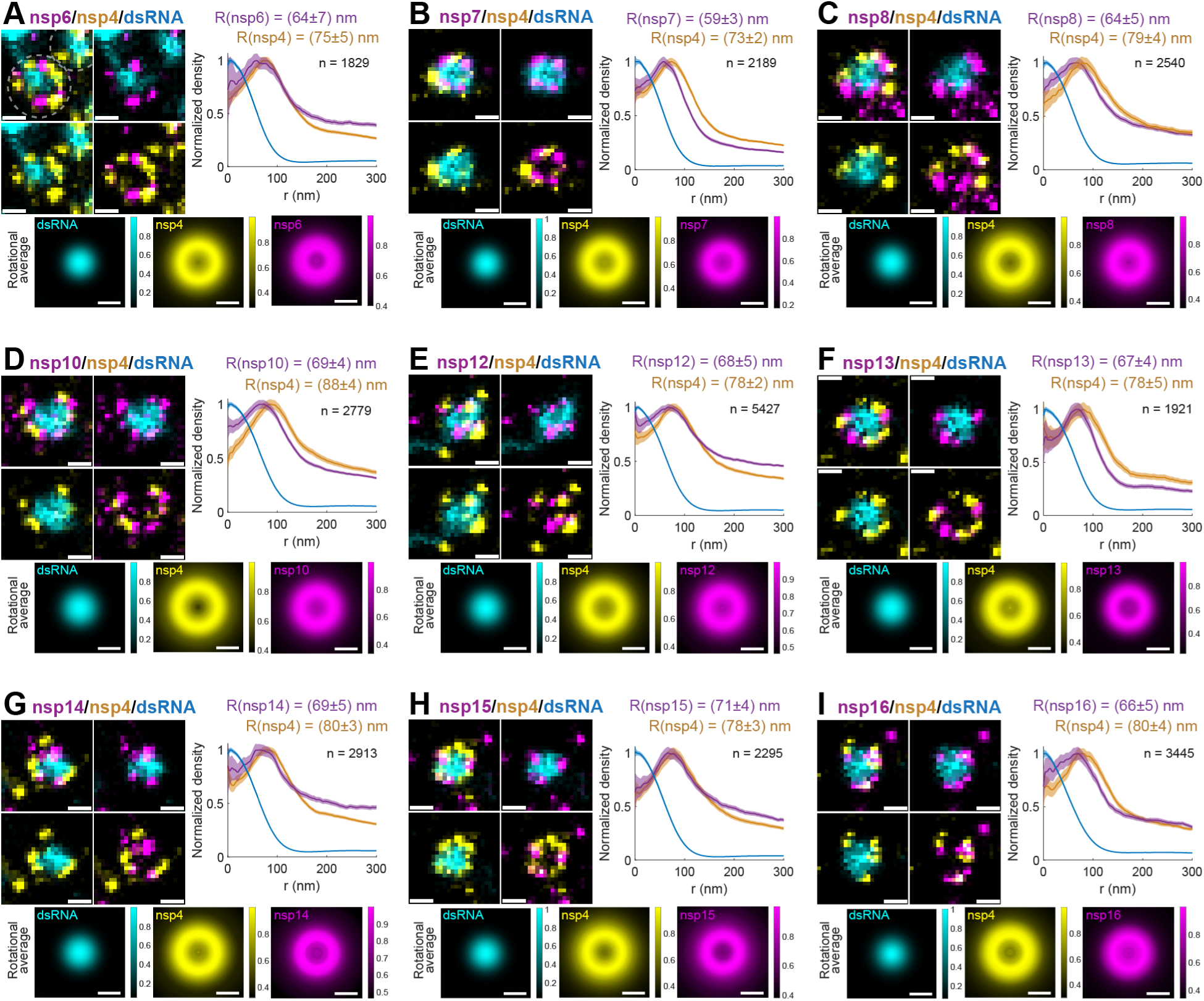
Nanoscale spatial organization of non-structural proteins within DMVs based on three-color SR imaging. (**A**–**I**) SR imaging and quantitative radial analysis maps the distribution of nsp6-nsp8 (**A**–**C**), nsp10 (**D**), and nsp12–nsp16 (**E**–**I**) multiplexed with nsp4 and dsRNA. Each panel displays a representative SR image of an individual DMV, the corresponding radial density distribution function *g*(*r*), and a rotationally averaged projection of the radial distribution. Dashed circles in **A** delineate two distinct DMVs. Peak radial positions (*R*) are indicated above the plots as the mean ± 95% confidence interval (CI), derived from Gaussian fitting of the peak region of *g*(*r*) following bootstrap resampling. The sample size (*n*) denotes the total number of DMVs analyzed per target combination. Analyses were performed on DMVs across 22 (**A**), 16 (**B**–**D**, **H**), 35 (**E**), 12 (**F**, **G**), and 18 (**I**) distinct early-stage cells. Scale bars: 100 nm.

**Supplementary Fig. 5:**
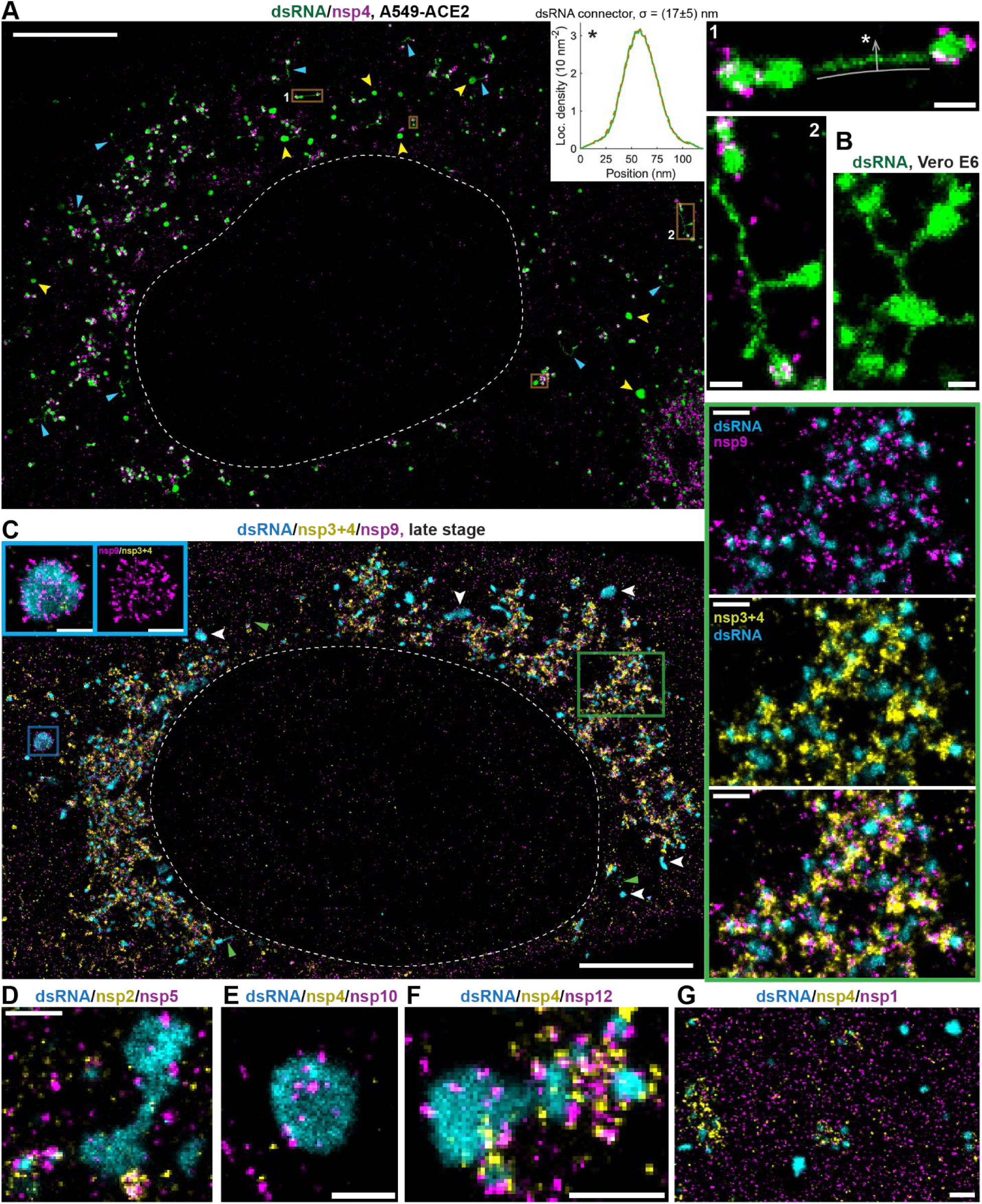
Subcellular organization of dsRNA connectors and granules in SARS-CoV-2–infected cells. (**A**) 2D dSTORM imaging of dsRNA and nsp4 reveals diverse dsRNA localization patterns. Magnified regions (right) corresponding to the yellow boxed areas in the main image highlight dsRNA connectors bridging DMVs. The inset displays a cross-sectional localization density profile of dsRNA, calculated orthogonal to the white curve in region 1, in the direction of the arrow (with longitudinal density averaged). The width of the fitted Gaussian function (*σ*, dashed line) is presented as the mean ± SD (n = 10 connectors from the represented cell). (**B**) Intercluster dsRNA connectors are similarly evident in Vero E6 cells labeled exclusively for dsRNA. (**C**–**G**) Three-color 2D SR images display large dsRNA granules and aggregates. (**C**) In late-stage infection, cells contain massive, round and elongated dsRNA structures. These aggregates lack DMV pore proteins (nsp3+4, labeled with a mixture of anti-nsp3 and anti-nsp4 primary antibodies) but recruit nsp9. Magnified insets (blue frame) detail a single large dsRNA aggregate devoid of pores. Conversely, the right panels (green frame) illustrate the typical perinuclear accumulation of dsRNA, nsp9 and nsp3+4 characteristic of late-stage DMVs and vesicle packets^5^. In panels **A** and **C**, arrowheads denote additional large dsRNA aggregates, while triangles highlight fiber-like dsRNA connectors. (**D**) An irregularly shaped dsRNA aggregate is decorated with discrete nsp5 puncta but lacks nsp2. (**E**, **F**) Large dsRNA structures largely devoid of nsp4 are nevertheless decorated with nsp10 (**E**) and nsp12 (**F**). (**G**) Voluminous, nsp4-deficient dsRNA granules occupy cytoplasmic zones entirely lacking nsp1. Images in **A** and **C**–**G** were acquired in A549-ACE2 cells. White dashed curves approximate the position of the nuclear envelope. Scale bars: 5 µm (main images in **A**, **C**), 500 nm (**D**–**G** and magnified boxes in **C**), and 200 nm (**B** and magnified boxes in **A**).

**Supplementary Fig. 6:**
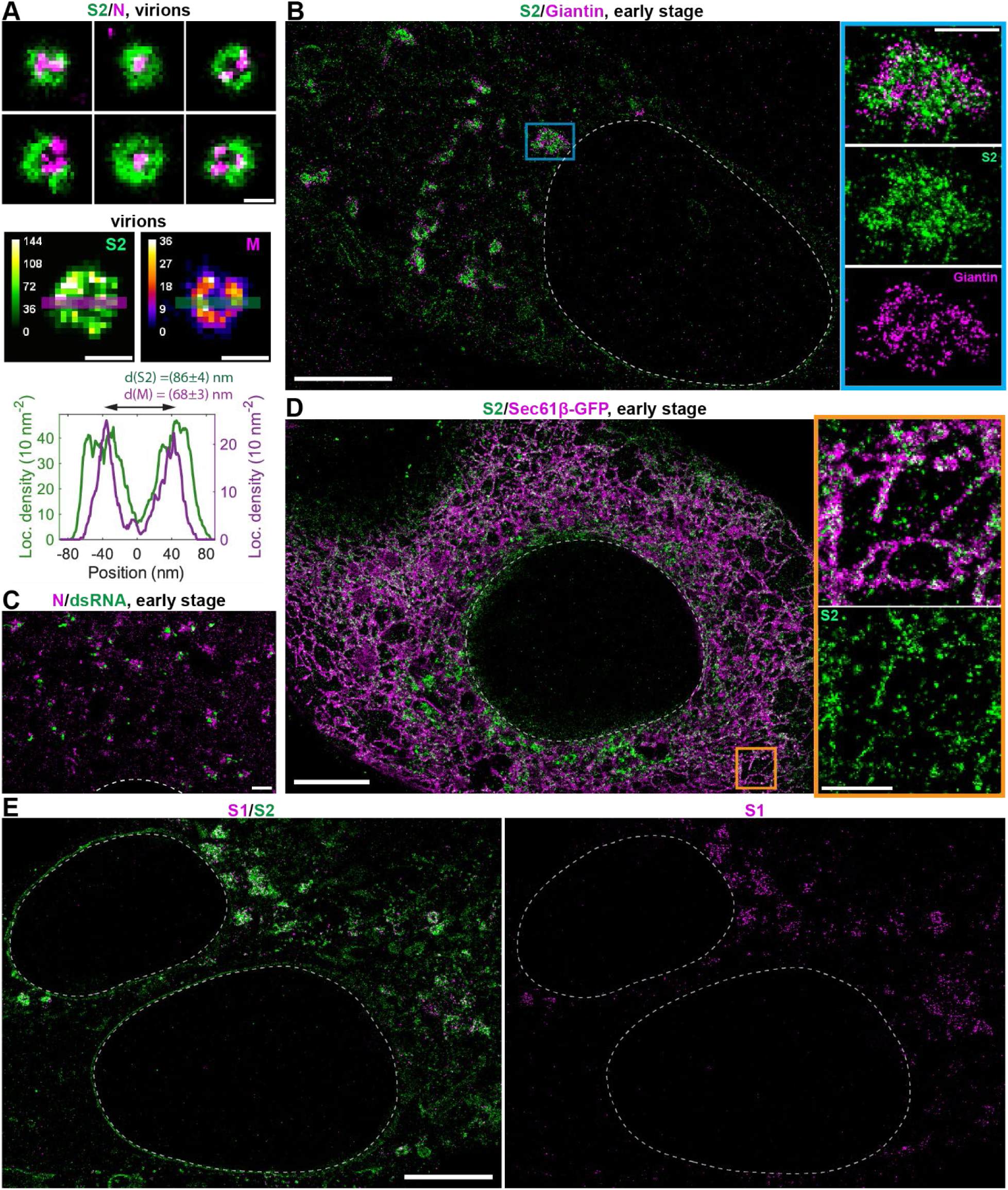
Subcellular localization and nanoscale organization of structural proteins in SARS-CoV-2–infected cells. (**A**) SR imaging of SARS-CoV-2 virions reveals the concentric organization of the S2 subunit around the N protein-labeled core (top). Both S2 and the membrane (M) protein form ring-like structures within the virions (middle). Cross-sectional localization density profiles, calculated along the indicated lines in the middle panels, demonstrate that M rings have a distinctly smaller diameter than S2 rings. The peak-to-peak distance between the two density lobes (*d*) was calculated by fitting the profiles with a sum of two Gaussians (*n* = 24 and 27 virions for M and S2, respectively). Values of *d* are represented as the mean ± 95% CI. Images of S2 and N were acquired from virions deposited on coverslips as previously described^5^, whereas M images were acquired from virions located at the cell periphery. (**B**) S2 localizes within Golgi bodies as indicated by the Giantin-labeled rim. (**C**) The N protein is distributed within the cytosol in close proximity to dsRNA-labeled ROs. (**D**) A fraction of S2 localizes to ER tubules (labeled with an anti-GFP antibody). Magnified regions (right) correspond to the orange boxed area in the whole-cell image. (**E**) Unlike S2, S1 subunit does not localize to either ER tubules or the nuclear envelope. Images were acquired in A549-ACE2 cells (**A**–**C**, **E**) and Vero E6-Sec61β-GFP cells (**D**). White dashed curves indicate the approximate position of the nuclear envelope. Scale bars: 100 nm (**A**), 1 µm (**C** and magnified images in **B**, **D**), and 5 µm (**E**, whole-cell images in **B**, **D**)

**Supplementary Fig. 7:**
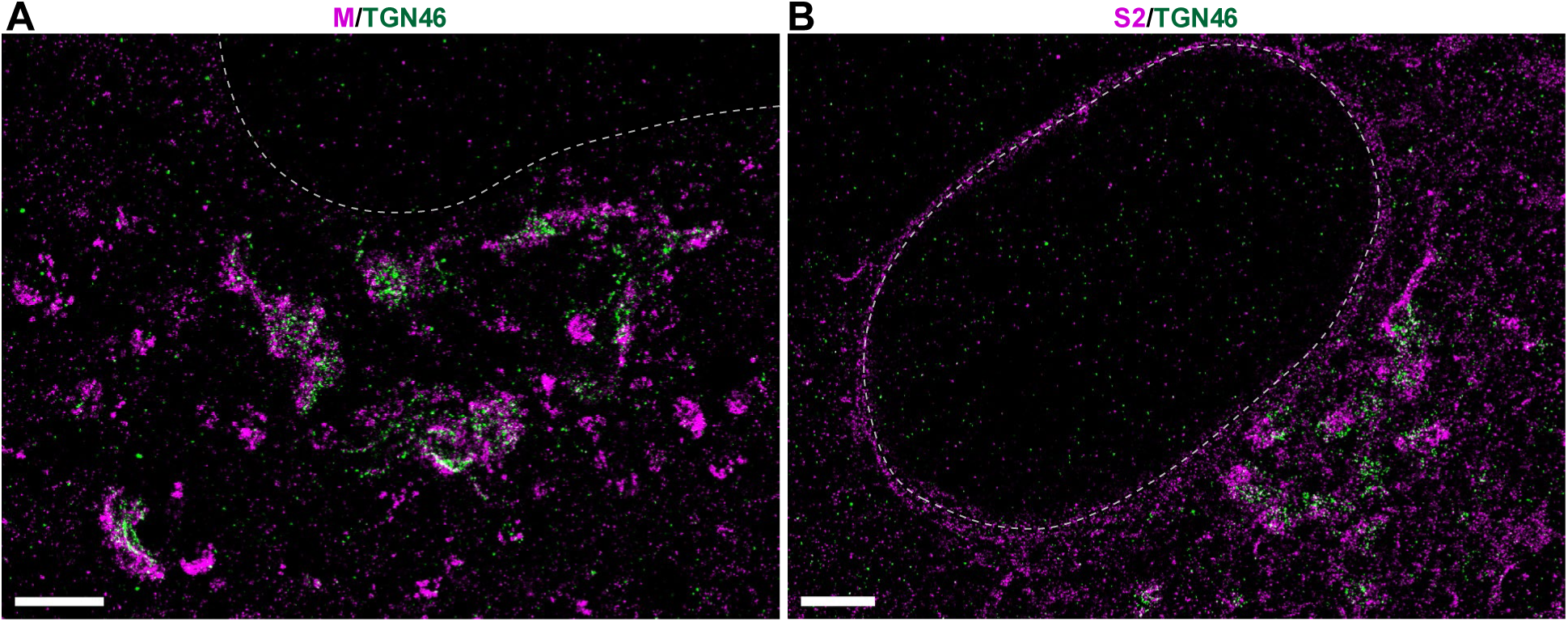
Structural proteins M and S2 localize to the trans-Golgi network. (**A**) Membrane protein (M) forms dense aggregates that are spatially juxtaposed to and partially overlap with TGN46 structures. (**B**) Similarly, spike subunit 2 (S2) clusters are observed adjacent to TGN46. White dashed curves indicate the approximate position of the nuclear envelope. Scale bars, 2 µm.

**Supplementary Fig. 8:**
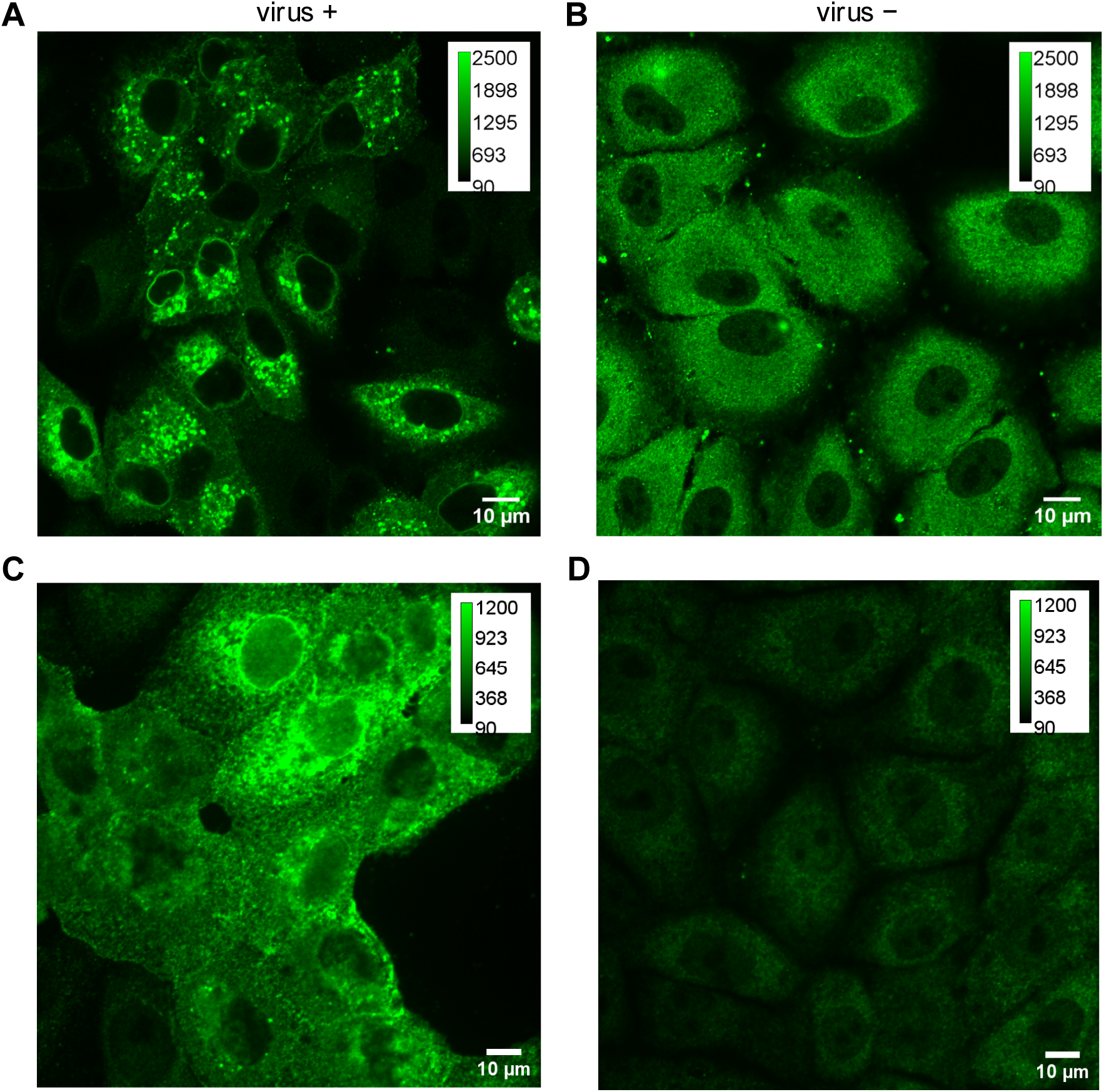
Validation of anti-S2 antibody specificity. (**A**–**D**) To confirm the specificity of the anti-S2 antibody and validate the genuine localization of the S2 subunit, confocal imaging was performed on SARS-CoV-2–infected (**A**, **C**) and uninfected control (**B**, **D**) cells. Distinct, specific localization of S2 to the nuclear envelope and endoplasmic reticulum (ER) is observed exclusively in the infected cells (**A**, **C**). Of note, the non-specific background signal in the uninfected control samples (**B**) is frequently higher than the signal observed in uninfected bystander cells within the infected samples (**A**). This is likely due to the depletion of the primary antibody by the highly abundant S2 antigen present in the infected cells. Images were acquired in A549-ACE2 cells (**A**, **B**) and Vero E6 cells (**C**, **D**), representing two biologically independent experiments. Scale bars: 10 µm.

**Supplementary Fig. 9:**
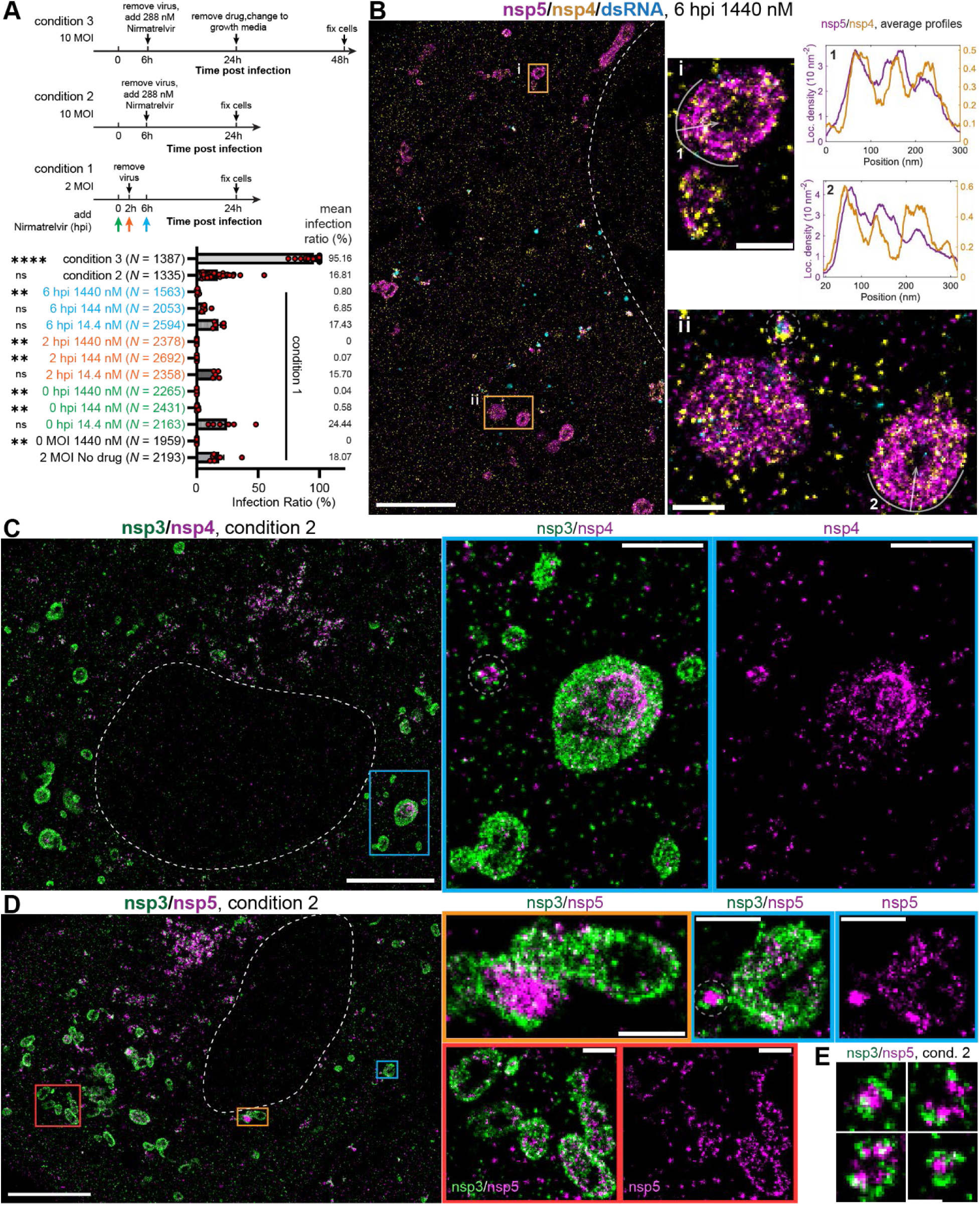
Nanoscale architecture of multilayered bodies in nirmatrelvir-treated, SARS-CoV-2–infected cells. (**A**) Schematics of the experimental timelines for conditions 1–3 (top) and a corresponding scatter dot plot of the infection ratio (infected cells / total cells) across these nirmatrelvir treatments (bottom). Individual data points represent the infection ratio per field of view. Statistical comparisons against the no-drug control were performed using a one-way ANOVA with Dunnett’s post hoc test. ns, not significant (p > 0.05); **, p < 0.01; ****, p < 0.0001. The sample size (*N*) denotes the total number of cells analyzed per condition. (**B**) SR imaging of a cell infected at an MOI of 2, treated with 1440 nM nirmatrelvir from 6 to 24 hpi, and fixed at 24 hpi. Magnified insets (center) highlight two regions containing large multilayered bodies (MLBs). Cross-sectional localization density profiles of nsp4 and nsp5 (right), calculated orthogonal to the white curves in the central insets (with longitudinal density averaged), demonstrate that both proteins exhibit a periodic, layered organization with a spacing of approximately 80 nm. (**C**–**E**) SR imaging of cells infected at an MOI of 10, treated with 288 nM nirmatrelvir from 6 to 24 hpi, and fixed at 24 hpi. Both nsp3/nsp4 (**C**) and nsp3/nsp5 (**D**) robustly localize to MLBs. Magnified colored boxes detail the diverse morphological variations of these MLBs. Morphologically typical DMVs are presented in **E**, with further instances delineated by dashed rings throughout panels (**B**–**D**). Experiments were performed in A549-ACE2 cells. White dashed curves indicate the approximate position of the nuclear envelope. Scale bars: 5 µm (main images in **B**–**D**); 1 µm (magnified boxes in **C**); 500 nm (magnified boxes in **B**, **D**); 200 nm (**E**).

**Supplementary Fig. 10.**
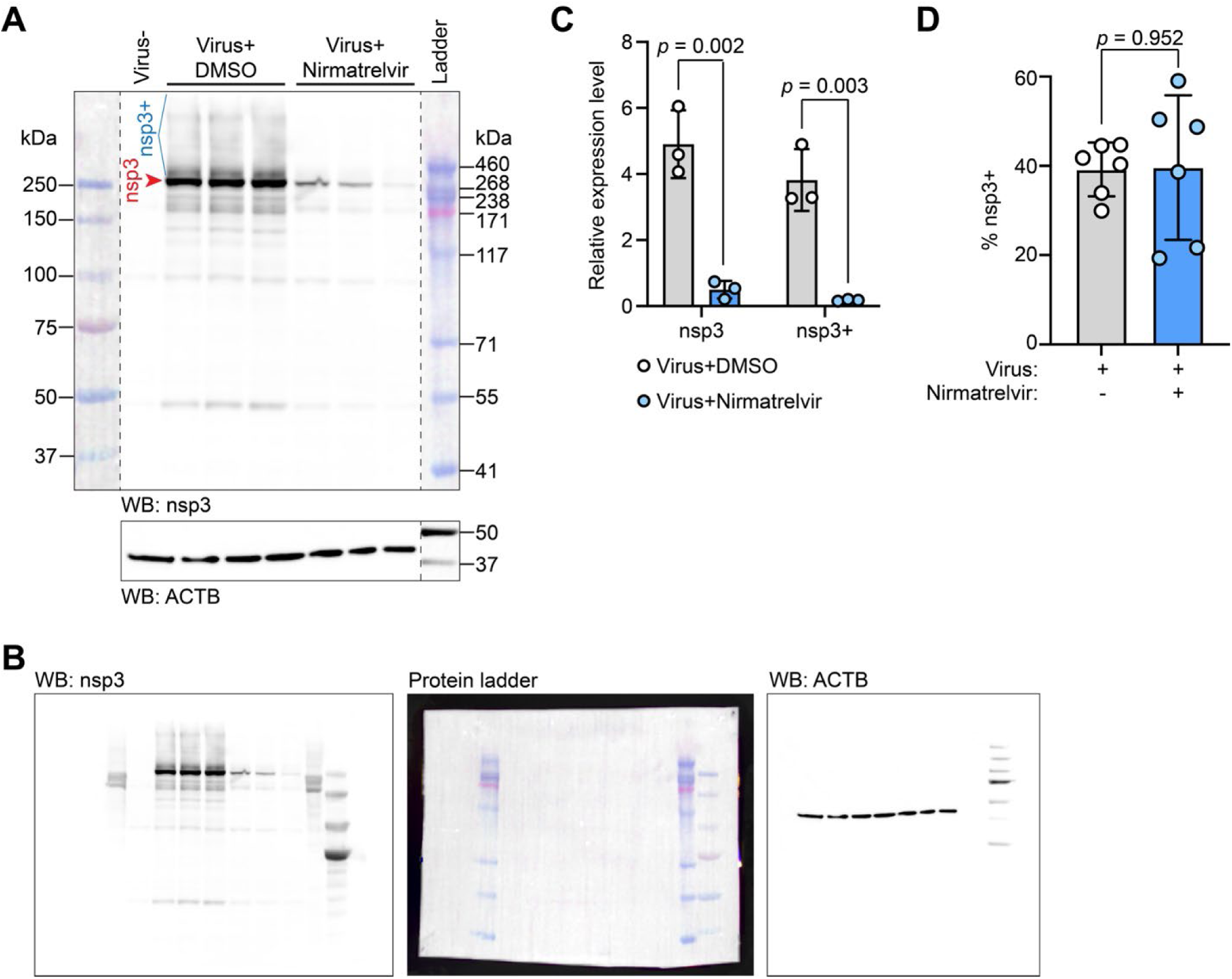
Western blot analysis of nsp3 cleavage under nirmatrelvir treatment. (**A**) Western blot analysis of nsp3 and ACTB proteins in cell lysates of A549-ACE2 cells without virus infection (Virus-), with SARS-CoV-2 virus infection and DMSO treatment (Virus+DMSO), and with SARS-CoV-2 virus infection and nirmatrelvir treatment (Virus+Nirmatrelvir). Overnight membrane transfer was conducted to guarantee large proteins were successfully transferred onto the membrane, as indicated by the 460 kDa protein ladder. Cleaved nsp3 protein bands are indicated by the red arrow. Uncleaved nsp3 protein bands (nsp3+) are indicated by the blue bracket. (**B**) Raw membrane images for the Western blot results in (**A**). Left: nsp3 blot captured using a gel imaging system. Middle: a direct photograph of the nsp3 blot membrane, taken with a smartphone, to clearly show protein ladders. Right: the corresponding ACTB loading control blot, captured using a gel imaging system. (**C**) Quantification of the relative expression level of nsp3 and nsp3+ proteins based on the Western blot result in (**A**). ACTB protein was used as a loading control for normalization. Three biological replicates. Two-sided, unpaired Student’s t-test. (**D**) Quantification of the percentage of nsp3+ proteins relative to total nsp3 proteins based on the Western blot result of (**A**). Three biological replicates and two experimental replicates. Two-sided, unpaired Student’s t-test.

Table 1 and Table 2 are provided as separate files.

